# Phylogenomics of *Aristolochia* subg. *Siphisi*a (Aristolochiaceae) reveals widespread incomplete lineage sorting and supports a novel pollinator-filtering hypothesis

**DOI:** 10.1101/2025.05.29.656634

**Authors:** Yifan Wang, Shuai Liao, Zirui Guo, Pan Li, Yusong Huang, Joyce G. Onyenedum

## Abstract

*Aristolochia* subgenus *Siphisia* constitutes a monophyletic lineage of predominantly lianescent species, with occasional shrubs or herbs, and is characterized by remarkable diversity in perianth morphology. Members of *Siphisia* serve as larval hosts for endangered Lepidoptera and are widely used in traditional medicine. Despite its ecological and ethnobotanical significance, *Siphisia* systematics remains unresolved due to limited genomic resources and insufficient phylogenetic signal across previously sampled loci. Here, we present a phylogenomically informed framework for *Siphisia*, integrating 46 newly collected accessions and seven public datasets across 44 taxa. Using genome skimming (∼30× coverage) and HybPiper, we recovered Angiosperms353 nuclear loci, including supercontigs, for phylogenetic reconstruction via concatenation and coalescent approaches. The resulting species trees resolve seven strongly supported monophyletic clades, each defined by distinct biogeographic patterns and morphological synapomorphies. Comparative plastome analyses from *de novo* assemblies explored quadripartite structure, plastid phylogeny, GC content, and gene synteny. Cytonuclear discordance was concentrated in species-rich Asian clades, while hybridization signals were rare and limited to deep backbone nodes in North American lineages. These results indicate that incomplete lineage sorting, rather than introgression, accounts for most gene–species tree conflicts and likely reflects strong reproductive isolation following speciation. We also revise a previously taxonomically ambiguous complex—the now well-supported *A. versicolor* species group—based on integrated phylogenomic and morphological evidence. Within this group, we describe five previously unrecognized cryptic species and identify a novel pollination syndrome involving floral adaptations for pollinator filtering. This syndrome may contribute to prezygotic isolation and recent diversification, and it challenges the prevailing assumption that *Aristolochia* pollination is universally governed by a trapping–release mechanism—suggesting this model may not apply to subg. *Siphisia*.

## 1. Introduction

*Aristolochia* L. is the largest and most diverse genus in Aristolochiaceae, comprised of over 600 species across a cosmopolitan distribution (Neinhuis et al., 2005; Ohi-Toma et al., 2006; Wagner et al., 2012). This genus exhibits remarkable biodiversity, particularly in its specialized perianth structures (González and Stevenson, 2000a, 2000b), unique pollination mechanisms (Alpuente et al., 2023; Oelschlägel et al., 2015, 2009; Rupp et al., 2021), and ecological significance as host plants for many Papilionidae Latreille species (Allio et al., 2021; Cini et al., 2019; Jain et al., 2021; Nishida and Fukami, 1989). As a member of the early diverging magnoliids, *Aristolochia* is taxonomically complex, housing three monophyletic subgenera: *Aristolochia*, *Pararistolochia* (Hutch. & Dalziel) Schimdt, and *Siphisia* (Duch.) Schimdt (Buchwalder et al., 2014; Neinhuis et al., 2005; Ohi-Toma et al., 2006; Ohi-Toma and Murata, 2016; Zhu et al., 2019b). Among these, subgenus *Siphisia* is the second-largest in terms of species richness and is currently undergoing rapid taxonomic expansion, with numerous newly discovered and described species—several of which have been assessed as endangered or even critically endangered upon publication. (Cai et al., 2020a, 2020b; Do et al., 2014, 2021a, 2021b, 2023; Do and Hoang, 2022; Gong et al., 2018; Guo et al., 2025; Huang et al., 2022; Kashung et al., 2022; Li et al., 2019; Lu et al., 2022; Luo et al., 2020; Ma et al., 2023; Nguyen et al., 2022; Ohi-Toma et al., 2021; Peng et al., 2019; Phan et al., 2021; Wang J. et al., 2020, 2021; Wang Y.F. et al., 2025; Watanabe-Toma et al., 2021; Xu et al., 2023; Yang et al., 2024; Zhang et al., 2024; Zhou et al., 2019; Zhu et al., 2017a, 2017b, 2018, 2019a, 2019b, 2019c, 2019d; Zhu and Ma, 2022). Many species of subgenus *Siphisia* are widely used in traditional Chinese medicine and other cultural ethnobotanical practices. (Bai et al., 2023; Poon et al., 2013; Shkryl et al., 2023), despite containing aristolochic acids with varying pathogenic and toxic effects among taxa (Allio et al., 2021; Das et al., 2022; Schmeiser et al., 2009; Shi et al., 2024). Given the continuously unfolding taxonomy of this subgenus and its foreseeable importance across domains such as traditional medicine and butterfly breeding, a well-resolved, species-level phylogeny for *Siphisia* is urgently needed. Such a framework is essential to support downstream efforts in taxonomy, conservation, pharmacological evaluation, and ecological research. Characterized by its unusually elaborate perianth morphology, *Siphisia* also holds considerable evolutionary significance for understanding patterns of floral diversification and adaptation. Despite its importance, species-level identification remains challenging due to elusive phenological traits such as cauliflory and irregular flowering cycles (Cai et al., 2024; Guo et al., 2025; Luo et al., 2020; Wang et al., 2025; Xu et al., 2023; Zhang et al., 2024), which complicate field recognition, hinder voucher identification and sampling, and ultimately impede taxonomic consistency across the literature. To date, no comprehensive systematic framework is available, and existing treatments remain preliminary and fragmented.

*Aristolochia* is well known for its distinctive floral morphology, hence the common name “pipevine.” As one of its three subgenera, *Siphisia* shares this architectural foundation, yet exhibits remarkable internal variation, with structurally divergent and highly specialized floral forms across the lineage. This striking morphological diversity suggests the possibility of varied or alternative reproductive strategies within *Siphisia* itself. However, despite the elaboration and distinctiveness of its floral traits, many *Siphisia* species have only recently been taxonomically validated, and their pollination biology remains poorly understood. In contrast, its sister subgenus, *Aristolochia* s.s. (i.e., subgenus *Aristolochia*), has a well-established reproductive paradigm supported by numerous species-specific studies (Alpuente et al., 2023; Berjano et al., 2009; Blatrix et al., 2024; Burgess et al., 2004; Nakonechnaya et al., 2024; Oelschlägel et al., 2015; Park and Kim, 2023; Rulik et al., 2008; Rupp et al., 2021; Trujillo and Sérsic, 2006). A hallmark strategy in this clade is the trapping–release mechanism, in which downward-pointing trichomes within the calyx tube act as physical barriers, temporarily confining pollinators until pollen release (Oelschlägel et al., 2009). Despite their phylogenetic proximity, the floral morphology of *Siphisia* is consistently and diagnostically different from that of *Aristolochia* s.s. (González and Stevenson, 2000b; Zhu et al., 2019b), raising the question of whether the canonical reproductive model applies to *Siphisia* at all. Could this subgenus have evolved an entirely distinct strategy—one equally complex in perianth architecture, but functionally different? If so, what selective pressures may have driven the evolution of such complexity in the absence of the same trapping mechanism? While these questions remain open, González and Pabón-Mora (2015) have emphasized the need for dedicated study of *Siphisia* pollination biology. Supporting this, Erbar et al. (2017) demonstrated substantial differences in nectary structure and secretion patterns between *Siphisia* and *Aristolochia* s.s., further suggesting the presence of a new reproductive system in this lineage.

While it is easy to be captivated by *Siphisia*’s most conspicuous flowers, our extensive sampling and fieldwork have also revealed intriguing variation in vegetative morphology, often associated with particular taxonomic groups. For example, several southern Chinese taxa appear to have evolved tuberous roots ranging from fusiform to moniliform, visually analogous to sweet potatoes (Guo et al., 2025; Wang et al., 2025; Xu et al., 2023). In terms of growth habit, some American species deviate from the predominant lianoid form that characterizes most of the lineage, instead exhibiting herbaceous or subshrub-like structures (Wagner et al., 2012). These observations suggest a complex pattern of infra-generic morphological evolution that remains largely untested within a systematic framework. One of the major limitations in previous studies of *Aristolochia* systematics, especially within *Siphisia*, has been poor taxonomic and geographic sampling. Many species are narrowly endemic and known from only a few individuals (Bai et al., 2023; Wanke et al., 2017), presenting considerable challenges in material acquisition, let alone the construction of comprehensive molecular datasets. When available, genetic data have often been limited to one or a few loci, which may resolve deep nodes but offer insufficient resolution at the species level (Li, 2019; Neinhuis et al., 2005; Zhu et al., 2019b). Fortunately, the field is now primed for progress. According to Guo et al. (2025), *Aristolochia* subg. *Siphisia* comprises over 120 species—a major expansion in our taxonomic understanding. With access to broad living collections and the increasing accessibility of next-generation sequencing (Ekblom and Galindo, 2011; Emelianova et al., 2023), we are now in a position to apply genome-wide data and advanced bioinformatics pipelines (Johnson et al., 2016; Leebens-Mack et al., 2019; Zhang et al., 2018) to establish a robust phylogenomic framework for the subgenus.

In this study, we aim to (1) reconstruct a broadly sampled phylogeny of *Aristolochia* subg. *Siphisia*, incorporating representatives from across its known biogeographic range and all major taxonomic sections, using genome-wide nuclear loci, and to propose a preliminary infrageneric classification; and (2) integrate this systematic framework with macroevolutionary and ecological insights—particularly those related to reproductive strategies and pollination biology. By doing so, we seek to clarify species boundaries, resolve long-standing taxonomic discrepancies, and uncover synapomorphies underlying *Siphisia* systematics.

## 2. Materials and methods

### 2.1. Taxon sampling, DNA extraction, sequencing

#### 2.1.1. Taxon sampling

Our sampling includes 44 species (53 accessions), with the majority (39 of 44) representing *Aristolochia* subg. *Siphisia*, including five newly described species (10 accessions). Among these, seven accessions were obtained from publicly available datasets in the NCBI Sequence Read Archive (Supplementary Table S1), while the remaining 46 were newly selected for this study. Five taxa from Aristolochiaceae were included as outgroups, following Wanke et al. (2021): *Aristolochia ringens* Vahl, *Aristolochia fimbriata* Cham., *Thottea hainanensis* (Merr. & Chun) Ding Hou, *Saruma henryi* Oliv., and *Lactoris fernandeziana* Phil. To ensure accurate species identification, all newly sequenced *Siphisia* samples were derived from living collections that had flowered for at least two consecutive years, allowing confirmation based on both reproductive and vegetative characteristics. Samples were collected through fieldwork or acquired via material exchange with botanical gardens between 2022 and 2024. Detailed information for each accession is provided in Supplementary Table S1.

#### 2.1.2. DNA extraction and sequencing

For newly sequenced samples, young leaves were selected in the field and immediately dried in silica gel at room temperature (22–28 °C) for 72 hours in preparation for DNA extractions.

Next-generation sequencing (NGS) data were generated across three sequencing batches processed by different providers. A total of 22 samples (Supplementary Table S1) were sequenced in April 2024 under project code BMK240417-BZ227-ZX01-01 by Beijing Biomarker Technologies Co., Ltd., with a target of at least 30 Gb of paired-end sequencing data per sample. Genomic DNA was extracted using the CTAB method (Doyle and Doyle, 1987), and quality was assessed using a Qubit™ 3.0 Fluorometer (dsDNA HS Assay Kit, Invitrogen) and 1% agarose gel electrophoresis. Only samples with DNA concentrations ≥5 ng/μL, total DNA ≥0.5 μg, and a main smear band ≥5 kb without visible degradation were retained for library preparation. Libraries were constructed using the VAHTS® Universal Plus DNA Library Prep Kit (ND617), and sequencing was performed on the Illumina NovaSeq 6000 platform (PE150 mode).

An additional 19 samples (Supplementary Table S1) were sequenced by Novogene Co., Ltd. (Beijing), with a minimum yield of 20 Gb of raw paired-end reads per sample, under projects X101SC23062418-Z01-J010_01 and X101SC23062418-Z01-J010_02. Genomic DNA was fragmented to an average size of 350 bp using a Covaris LE220R-plus ultrasonicator, followed by end-repair, A-tailing, adapter ligation, PCR amplification, and purification using AMPure XP beads. Library quality was evaluated using an Agilent 5400 (AATI), and quantification was performed using real-time PCR (1.5 nM). Sequencing was conducted on the DNBSEQ-T7 platform. For both batches (totaling 41 samples), sequencing depth was estimated to exceed 30× genome coverage, sufficient for plastome assembly and nuclear gene recovery, based on available genome size estimates for *Aristolochia fimbriata* (257.7 Mb), *Aristolochia contorta* Bunge (209.3 Mb), *Aristolochia californica* Torr. (661 Mb), and *Aristolochia manshuriensis* Kom. (525 Mb) (Chaturvedi et al., 2024; Cui et al., 2022; Hu et al., 2025; Qin et al., 2021).

Three additional samples (Supplementary Table S1; sampleSource: LI_ZJU) were sequenced by the China National GeneBank (CNGB) with 6 Gb of paired-end data per sample. Two additional samples (sampleSource: ZUO_XTBG) were sequenced at XTBG.

The sequencing protocol manuals are under contractual restriction with the service providers and cannot be publicly distributed, but complete methodological details are available upon request for research purposes. All sequencing data have been deposited in the NCBI Sequence Read Archive under accession number PRJNA1242357 (Supplementary Table S1).

### 2.2. Nuclear and plastome dataset construction

#### 2.2.1. Nuclear locus recovery

All newly generated and open-source SRA datasets consisted of paired-end short-read FASTQ files. A standardized pipeline was applied to recover genomic loci suitable for phylogenomic analyses, using the Angiosperms353 probe set (Johnson et al., 2019) as the reference. To construct the reference dataset, target gene sequences were retrieved from the Royal Botanic Gardens, Kew Tree of Life Explorer (https://treeoflife.kew.org/) (Baker et al., 2022). Within *Aristolochia*, the only available Angiosperms353 reference was derived from *Aristolochia littoralis* Parodi, originally sourced from the One Thousand Plant (OneKP) Transcriptomes Initiative (Leebens-Mack et al., 2019). This dataset contained 330 of the 353 target loci, totaling 234,198 bp.

Raw reads from 53 accessions were quality-filtered using Trimmomatic v0.39 (Bolger et al., 2014) on the NYU IT High Performance Computing (HPC) cluster. Reads were processed in paired-end mode with adapter removal, and quality trimming was conducted using a 4-base sliding window (minimum average Phred score 15), with reads below 36 bp discarded. A Phred score threshold of 33 was applied to ensure high-confidence base calls.

Trimmed reads were processed with HybPiper v1.3.1 (Johnson et al., 2016) to extract target loci in DNA mode, using *A. littoralis* as the reference and bwa (Li and Durbin, 2009) as the aligner. HybPiper automatically flagged genes with anomalously high read depth suggestive of paralogs, which were excluded to ensure accurate ortholog recovery. Following gene recovery, we applied HybPiper’s built-in retrieve_sequences.py script to reconstruct supercontigs (exon + intron) for each sample. Note that while *A. littoralis* was used as a reference for target mapping, it was not included in downstream phylogenomic analyses due to its exon-only nature.

#### 2.2.2. Plastome assembly, annotation and structural analysis

Plastome assembly was performed directly from raw short-read datasets following the streamlined protocol proposed by Jost and Wanke (2024). *De novo* assembly was conducted using GetOrganelle v1.7.5 (Jin et al., 2020), which employs a seed-and-extend algorithm optimized for reconstructing circularized chloroplast genomes. Assemblies were executed with the following parameters: --fast -k 21,77,127 -F embplant_pt, using a custom seed file from the *Aristolochia kwangsiensis* Chun & F.C. How ex C.Y. Cheng plastome (GenBank Accession: OP950693.1).

Assembly results yielded fully circularized plastomes in 49 accessions, while four accessions produced high-coverage linear scaffolds (Supplementary Table S1). For samples that failed to generate circular genomes, scaffolded assemblies were imported into Geneious Prime 2025.0.3 for manual curation, including gap closure and circularization, strictly following the procedures described in Jost and Wanke (2024). Plastome annotation was conducted with reference to two representative Piperales plastid genomes: *Piper nigrum* L. (GenBank Accession: NC_034692) and *Thottea sumatrana* (Merr.) Ding Hou (GenBank Accession: NC_065017), to ensure consistency in gene boundary and structure annotation. Annotation and structural validation, including identification and confirmation of inverted repeat (IR) boundaries, were performed in Geneious Prime 2025.0.3, with manual correction as needed for gene boundary accuracy.

Chloroplast junction pattern visualization was carried out using CPJSdraw (Li et al., 2023). Final plastome visualization was rendered using the OGDraw online platform (https://chlorobox.mpimp-golm.mpg.de/OGDraw.html) (Greiner et al., 2019). GC content (total, regional, and codon-position-specific) was calculated from annotated GenBank files using a custom script developed in this study (available at https://github.com/Pipevine1122/A.versicolor under script name: gc_content_parser.py).

### 2.3. Phylogenomic reconstruction

#### 2.3.1. Nuclear phylogeny inference

For nuclear phylogenetic reconstruction, each recovered Angiosperms353 supercontig locus was aligned individually using MAFFT v7.475 (Katoh et al., 2002) with the L-INS-i strategy, suitable for highly accurate alignment of medium-sized datasets. All alignments were performed on the NYU IT High Performance Computing (HPC) system.

The resulting gene alignments were concatenated into a supermatrix using a custom Python script developed in this study (available at GitHub: https://github.com/Pipevine1122/A.versicolor; script name: 11.D.concatenate.py), ensuring consistent gene order across samples. A partition file recording the order and length of each locus was also generated. Phylogenetic inference of the concatenated matrix was conducted using IQ-TREE 2 (Minh et al., 2020), with the ModelFinder Plus (MFP) function to determine the optimal substitution model for each partition, and 2000 ultrafast bootstrap replicates to assess node support.

In parallel, single-gene trees were inferred for each aligned locus using IQ-TREE 2 under the same bootstrap settings. These individual gene trees were then used to construct a coalescent-based species tree with ASTRAL-III (Zhang et al., 2018). *Lactoris fernandeziana* was designated as the outgroup using the --outgroup option. The -t 2 flag in ASTRAL-III was applied to obtain local posterior probability (localPP) scores for internal branches (Shang et al., 2024), providing a probabilistic measure of node support based on quartet frequencies.

#### 2.3.2. Complete plastome phylogeny

For plastid-based phylogenetic reconstruction, we strictly followed the methodology outlined in Jost and Wanke (2024). Annotated protein-coding regions from each chloroplast genome were extracted using a custom script developed in this study (available at GitHub: https://github.com/Pipevine1122/A.versicolor; script name: 03.D.extract_genes.py). Each gene was aligned using MAFFT v7.475 (Katoh et al., 2002) with the L-INS-i strategy to maximize alignment accuracy.

A total of 340 aligned nuclear loci were included. For samples missing specific loci, the corresponding positions were retained in the alignment as question marks. The aligned regions consisted of fully recovered supercontigs—including both exons and introns—which were concatenated into a single supermatrix with preserved gene order. During concatenation, individual gene lengths were recorded and a corresponding partition file was generated using another custom script from this study. Maximum likelihood phylogenetic inference was performed using IQ-TREE 2 (Minh et al., 2020), applying the ModelFinder Plus (MFP) function to identify the best-fit substitution model for each partition. Bootstrap support values were estimated using 2,000 ultrafast bootstrap replicates.

To ensure both methodological and sampling consistency with the nuclear dataset, we did not include publicly available *Siphisia* plastomes. Instead, all taxa with previously published plastid genomes were resampled, resequenced, and assembled *de novo* under identical protocols to maintain uniformity in assembly and annotation quality, ensure taxon set congruence with the nuclear phylogeny, and avoid potential chimeric artifacts in downstream comparisons.

#### 2.3.3. Ancestral state reconstruction

We reconstructed ancestral states for discrete floral traits associated with the RICH syndrome in the A. versicolor species group using a custom R script (available at https://github.com/Pipevine1122/A.versicolor under script name: Ancestral_state.R). The analysis focused on ten taxa either exhibiting or phylogenetically proximate to the syndrome (Supplementary File S7, with one representative accession selected per species when multiple samples were available: *Aristolochia nitida* Y.Fan Wang, Z.R.Guo & J.G.Onyenedum (NCBI BioSample Accession: SAMN47587624), *Aristolochia magnopurpurea* Y.Fan Wang, Z.R.Guo & J.G.Onyenedum (NCBI BioSample Accession: SAMN47587622), and *Aristolochia yueguiensis* Y.Fan Wang, Z.R.Guo & J.G.Onyenedum (NCBI BioSample Accession: SAMN47587614).

Character states were compiled as a binary matrix (Supplementary File S7), coded as 1 (absent) or 2 (present). Trees were rooted with *Lactoris fernandeziana* and pruned to match taxa in each trait matrix. For each trait, we implemented ancestral state reconstruction on the concatenated supermatrix tree (Supplementary File S3) using stochastic character mapping, based on the transition rate matrix (Q) estimated from the best-fitting model among equal rates (ER), symmetrical (SYM), and all-rates-different (ARD) models. Model selection was performed via corrected Akaike Information Criterion (AICc). All reconstructions used 1,000 simulations under the selected model.

### 2.4. Test for hybridization/introgression and incomplete lineage sorting

#### 2.4.1. Node discordance assessment between gene and species trees

To evaluate discordance between single-gene trees and the coalescent-based species tree, we used the previously inferred gene trees alongside the ASTRAL-III species tree. Phyparts (Smith et al., 2015) was applied to quantify unique, conflicting, and concordant bipartitions, enabling assessment of gene tree–species tree incongruence at each node. Visualization of Phyparts results was conducted using a modified script from https://github.com/mossmatters/MJPythonNotebooks, allowing detailed node-by-node evaluation of topological concordance and conflict across the phylogeny.

#### 2.4.2. Reticulation phylogeny and hybridization

To assess potential reticulation and hybridization events, we subsetted nine representative species based on our phylogenomic results: *Aristolochia westlandii* Hemsl., *Aristolochia dabieshanensis* C.Y.Cheng & W.Yu, *Aristolochia griffithii* Hook.f. & Thomson ex Duch., *Aristolochia serpentaria* L., *Aristolochia tricaudata* Lem., *Aristolochia kunmingensis* C.Y.Cheng & J.S.Ma, *A. californica*, *A. manshuriensis*, and *Lactoris fernandeziana* (as outgroup). These species were chosen to represent the major clades identified in the ASTRAL-based nuclear phylogeny.

The same HybPiper-recovered datasets corresponding to these nine taxa were selected, and the ASTRAL-III pipeline was rerun to generate a coalescent-based species tree for this reduced dataset. To infer reticulation events and assess gene tree discordance potentially driven by hybridization, we applied the maximum pseudo-likelihood method (InferNetwork_MPL) implemented in PhyloNet (Than et al., 2008). Network inference was conducted by allowing a maximum of 1, 2, or 3 reticulation events. For each setting, 10 independent searches were performed to reduce the likelihood of convergence on local optima. The log-likelihood, Akaike Information Criterion (AIC), and Bayesian Information Criterion (BIC) were computed for each model to evaluate model fit and complexity. The resulting networks were visualized using Dendroscope 3 (Huson and Scornavacca, 2012).

#### 2.4.3. ILS detection

To further investigate the underlying causes of phylogenetic discordance, we used Phytop (Shang et al., 2024) to quantify the contributions of incomplete lineage sorting (ILS) and introgression/hybridization (IH, hybridization and introgression here are treated as a single combined factor in this context). The input species tree was the ASTRAL-inferred topology with branch support derived from quartet frequencies.

### 2.5. Morphological examination

Morphological examination was conducted using a Fujifilm X-T3 digital camera equipped with a Laowa 65mm f/2.8 Ultra Macro lens to document all freshly collected floral specimens. Flowers were immediately dissected upon collection to examine perianth adaxial structures. Dissected perianth tissues were subsequently pressed and dried as voucher specimens. Epidermal cells of the dried perianth were observed using an Olympus CK2 stereo macroscope under 20× magnification.

### 2.6. Conservation assessment

The conservation status was assessed following the IUCN Red List Categories and Criteria (2012) and the most recent Guidelines for Using the IUCN Red List Categories and Criteria (2024). The Extent of Occurrence (EOO) and Area of Occupancy (AOO) were calculated using GeoCAT (Geospatial Conservation Assessment Tool; Bachman et al., 2011), based on field observations, ecological data, and verified distribution records to estimate the species’ geographic range and assess potential threats.

## 3. Results

### 3.1. Phylogenomic analysis

#### 3.1.1. nuclear Phylogenomic topology of Siphisia

Phylogenomic tree reconstruction using Angiosperms353 supercontigs, through both concatenated (“supermatrix”) and coalescent-based (ASTRAL) approaches, consistently recovered subg. *Siphisia* as a monophyletic lineage within *Aristolochia* s.l. (Figure 1; Supplementary Figure S1), with full statistical support across both methods (LPP = 1.0; BS = 100). The combined nuclear phylogenies also supported *Aristolochia* s.l. as a monophyletic genus, with *Thottea* recovered as its closest outgroup. The complete chloroplast phylogeny (Figure 2; Supplementary Figure S2) recovered the same subgeneric relationships, corroborating the nuclear results and again resolving *Aristolochia* s.l. as a fully supported monophyletic group (BS = 100), with identical placement of outgroups.

**Figure 1.**
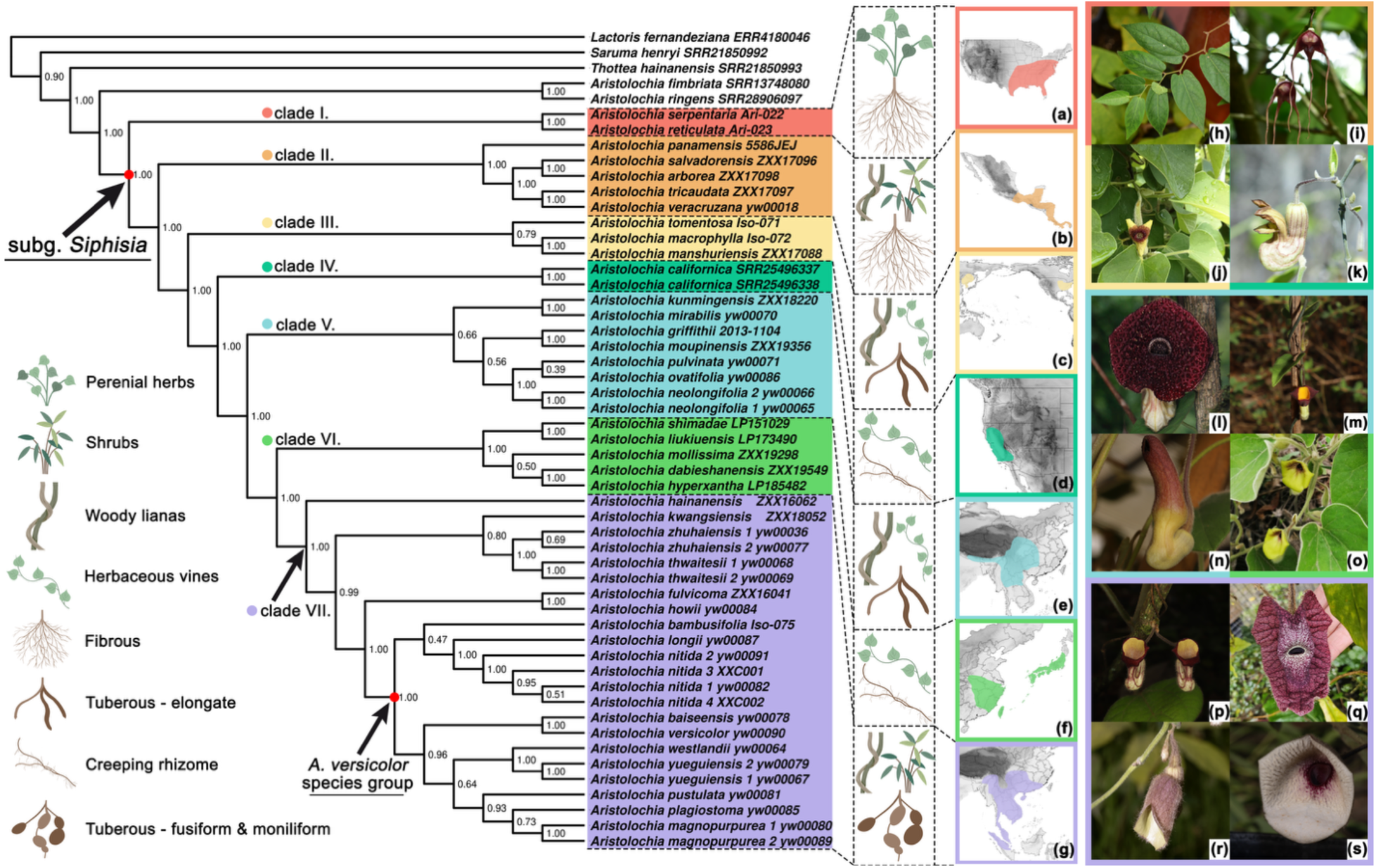
*Aristolochia* subg. *Siphisia* species tree inferred from Angiosperms353 supercontigs using ASTRAL-III, showing seven distinct monophyletic clades with corresponding synapomorphies in growth habit, root morphology, biogeographic distribution, and floral traits. **(a–g)** Geographic distribution maps of each clade: **(a)** Eastern and Southeastern United States; **(b)** Southern Central America from Panama to southern Mexico within the tropical rainforest zone; **(c)** Northeastern Asia (including Northeast China, Korea, and parts of Russia) and Eastern United States, suggesting possible species migration via the Bering land bridge; **(d)** California, along the western coast of the United States; **(e)** Southwestern China, northern Myanmar, northern Thailand, and northeastern India—forming a Hengduan Mountain–southern Himalayan biodiversity hotspot; **(f)** Eastern Sino-Japanese floristic region, reflecting climatic and biocommunity congruence; **(g)** Southern China, Indochinese Peninsula, Malay Peninsula, and the island of Sumatra, all within tropical monsoon or rainforest climatic zones. **(h–s)** Representative in situ images of species from each clade. Colored outer rims indicate their corresponding clades, matched in both color and label in the phylogenomic tree: **(h)** *A. serpentaria*; **(i)** *A. tricaudata*; **(j)** *A. tomentosa*; **(k)** *A. californica*; **(l)** *A. griffithii*; **(m)** *A. kunmingensis*; **(n)** *A. ovatifolia*; **(o)** *A. mollissima*; **(p)** *A. hainanensis*; **(q)** *A. westlandii*; **(r)** *A. plagiostoma*; **(s)** *A. zhuhaiensis*.

**Figure 2.**
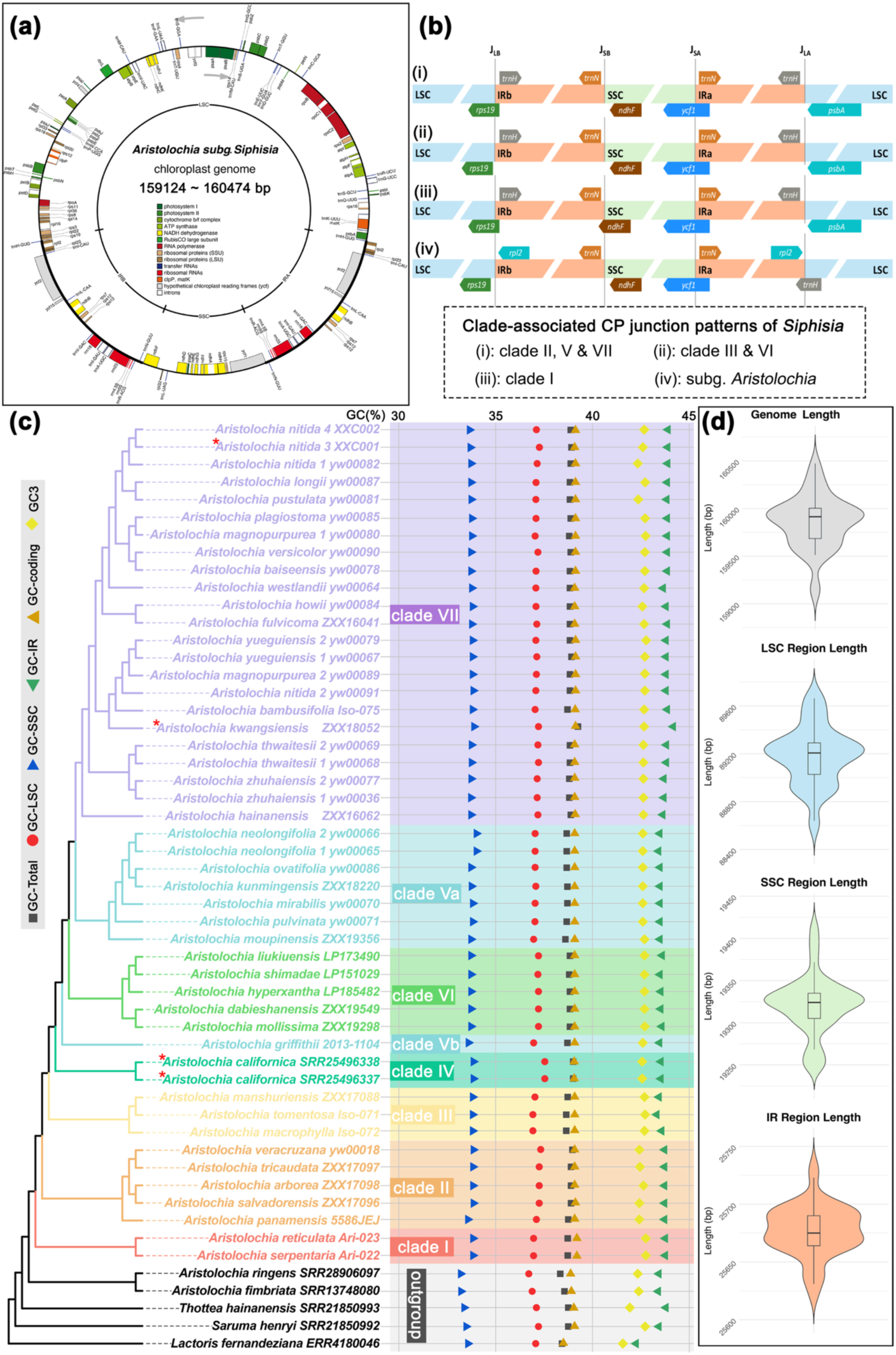
Comparative plastome analysis of *Aristolochia* subg. *Siphisia*. **(a)** Plastome synteny map of *Siphisia* samples automatically circularized using GetOrganelle. Only fully circularized samples are considered; manually circularized assemblies are excluded. All genomes exhibit a highly conserved structure with identical gene content and synteny across the subgenus. **(b)** Clade-associated patterns of chloroplast junctions in *Aristolochia* subg. *Siphisia*. Three distinct junction types are identified among the circularized plastomes. Patterns **(i)** and **(ii)** have been previously reported in *Siphisia* plastomes; pattern **(iii)** is newly documented here and is unique to the two herbaceous, non-twining U.S. endemics (*A. serpentaria* and *A. reticulata*), characterized by a novel placement of the *ndhF* gene extending from the SSC into the IRb region. This arrangement is reported for the first time in *Aristolochia* s.l. Clade assignments correspond to those defined by the nuclear coalescent phylogeny in the main text. **(iv)** Junction pattern observed in two *Aristolochia* s.s. species, included for comparison. **(c)** Maximum likelihood phylogeny based on complete chloroplast CDS and RNA genes, with 2,000 bootstrap replicates. Clade designations match those from nuclear phylogenomics. While overall plastome structure is conserved, topological conflict exists at both deep and shallow nodes, particularly between clades V and VI. For downstream analysis and discussion, *A. griffithii* is labeled as clade Vb, while all remaining taxa within Clade V are designated as Va. Middle scatter plots show GC content across genomic regions (GC-total, GC-LSC, GC-SSC, GC-IR, and GC3), indicating consistent values across *Siphisia* samples. Samples marked with red asterisks in phylogenetic tree failed automatic circularization via GetOrganelle despite multiple attempts and were later circularized manually. Clade colors are consistent with nuclear phylogenomic clade assignments and correspond to those in Figure 1. **(d)** Violin plots of plastome and regional genome lengths (LSC, SSC, IR) across all samples.

In the coalescent-based species tree, most nodes were recovered with very strong support (LPP = 1.0), particularly across the backbone topology (Figure 1). All bifurcation nodes among outgroup taxa also received high support (LPP ≥ 0.9). Within subg. *Siphisia*, the clade sister to the rest comprises *Aristolochia reticulata* Duch. and *A. serpentaria*, forming a monophyletic group with full support (LPP = 1.0). The second nested lineage is a fully supported clade (LPP = 1.0) composed of five species. The next monophyletic group includes three species—*A. macrophylla* Lam., *A. tomentosa* Sims, and *A. manshuriensis*. The subsequent clade is monotypic, consisting solely of *A. californica*. All remaining taxa are grouped into three major monophyletic clades. One well-supported clade includes eight taxa; another comprises five species; and the final clade, which is the most species-rich, contains the largest number of accessions (23). While most nodes exhibit full support, minor reductions in LPP (<0.9) were observed at several terminal tip bifurcating nodes, such as between two individuals of *Aristolochia zhuhaiensis* Y.Fan Wang & Z.R.Guo, which are morphologically identical. Overall, in the entire topology, 42 of 53 internal nodes (79.2%) received LPP ≥ 0.9, indicating strong support for the inferred relationships. Nodes with lower support were associated with shallow, tip-level divergences.

In the concatenated supermatrix phylogeny (Supplementary Figure S1), the overall backbone topology was highly congruent with that of the coalescent-based species tree, with all major clades showing identical taxon composition and the same sequential divergence pattern. Subg. *Siphisia* was consistently resolved as monophyletic and sister to *Aristolochia* s.s., with this pair forming a clade sister to *Thottea*. Although both phylogenies were inferred from the same nuclear loci, the concatenated tree yielded slightly higher overall support: 45 of 53 internal nodes (84.9%) received full bootstrap support (BS = 100), and only two nodes (3.7%) fell below BS = 90. Short internal branch lengths within *Siphisia* (Supplementary Figure S1, right panel) suggest recent diversification, whereas the relatively long branch leading to *A. californica* in both trees indicates deeper divergence. The most prominent topological conflicts were concentrated among the three deeply nested monophyletic groups, with several notable shifts in the placement of specific taxa between trees (Supplementary Figure S2).

#### 3.1.2. Systematics and Morphological Synapomorphies

The seven monophyletic clades, recovered consistently across both nuclear phylogenomic topologies, are further supported by distinct vegetative synapomorphies and biogeographical patterns. We define these preliminary clades as Clades I–VII and summarize their traits and systematic implications herein (Figure 1).

Clade I, occupying the earliest diverging position in both phylogenies, is morphologically and geographically distinct. It comprises two herbaceous North American species with fibrous root systems. Unlike the majority of congeners that are woody lianas, these are perennial herbs that sprout annually from fibrous roots. Both species are distributed in the eastern United States. Clade II includes five Central American endemics forming a monophyletic group ranging from woody lianas to subshrubs. These species inhabit rainforest environments and are reported to mimic fungal fruiting bodies both visually and chemically to attract and trap pollinators (Erbar et al., 2017; Wagner et al, 2012). Clade III consists of three species from eastern North America and northeastern Asia—herbaceous vines or woody lianas with elongated tuberous roots. This group reflects a classic eastern Asia–eastern North America disjunction, likely stemming from ancient migration via the Bering land bridge. *A. macrophylla* is found in Northeast Asia (China, the Korean Peninsula, and the Russian Far East), while *A. tomentosa* and *A. macrophylla* occur in eastern North America. These species share notable morphological similarities. Clade IV is a monotypic lineage formed by *A. californica*, endemic to coastal California. It is characterized by herbaceous vines with creeping underground stems that resprout annually. Its isolated phylogenetic position and geographic distribution, combined with a long branch length, suggest substantial genetic divergence from other *Siphisia* taxa (Figure S2). These four clades are composed almost of species from the Americas.

In contrast, the remaining three deeply nested clades, each containing numerous Asian endemic taxa, represent the majority of species diversity in the genus. Clade V is centered in southwestern Yunnan, particularly the Hengduan Mountains and the southern Himalayan biodiversity hotspot. In our study, it includes eight accessions representing seven species. Species in this clade possess yam-like tuberous roots that may extend underground for several meters, as observed in fieldwork. Some taxa within this group have evolved extremely enclosed perianths that restrict access to only minute pollinators (e.g., Figure 1n). However, floral morphology within this clade is highly variable, and no single defining synapomorphy can be easily extracted. Most species are vegetatively climbing or lianoid in habit, with occasional sympatry. Clade VI comprises species from eastern China and Japan, regions sharing parallel climatic and biogeographic histories that promote lineage divergence under similar ecological conditions. Unlike Clade V, species in Clade VI possess slender creeping rhizomes and remain herbaceous throughout life, lacking lignified or thickened stems. Clade VII, the most species-rich clade (16 taxa, 23 accessions), spans southern and southwestern China through the Indochina Peninsula and down to the Malay Peninsula and Sumatra. This lineage is characterized by unique fusiform to moniliform tuberous root systems not observed in other clades. These roots likely function to anchor the plants within crevices of karst limestone, environments with minimal humus layers. Most members are perennial woody lianas with robust, lignified stems, though two exceptions: *Aristolochia thwaitesii* Hook. and *A. zhuhaiensis*, are subshrubs.

These clade delineations will serve as the framework for subsequent analyses and discussions.

### 3.2. Comparative plastome genomics

We successfully recovered 49 complete circularized plastomes from 53 accessions (Supplementary File S1), including 46 newly sequenced samples and 7 from public SRA datasets. Four samples: *A. kwangsiensis* (ZXX18052), *A. nitida* sp. nov. (XXC0001), and two accessions of *A. californica* (SRR25496338, SRR25496337)—failed to generate fully circularized genomes and were manually curated (Supplementary Table S1).

All automatically circularized plastomes conformed to the canonical quadripartite structure, with total lengths ranging from 158,837 bp (*Aristolochia salvadorensis* Standl., ZXX17096) to 160,474 bp (*A. tomentosa*). The Large Single Copy (LSC) region ranged from 88,325–89,665 bp, the Small Single Copy (SSC) from 19,205–19,414 bp, and each of the Inverted Repeats (IRa and IRb)—which are identical in length—ranged from 25,621–25,733 bp. (Figure 2d). Gene annotation, following the workflow of (Jost and Wanke, 2024), revealed a conserved set of 114 unique genes across all *Siphisia* accessions: 80 protein-coding, 30 tRNA, and 4 rRNA genes. Gene synteny was identical across taxa (Figure 2a). Sixteen genes had one intron, while *clpP*, *ycf3*, and *rps12* contained two; *rps12* was trans-spliced, consistent with other Piperales plastomes.

Despite the conserved gene content, we observed structural variation at junctions between genome regions. Three distinct IR boundary patterns were detected (Figure 2b). Pattern (i), found in clades II, V, and VII, had *rps19* spanning the LSC-IRb junction and *ndhF* entirely within SSC. Pattern (ii), present in clades III and VI, placed *rps19* entirely within the LSC, consistent with prior reports (Bai et al., 2023; Jost and Wanke, 2024). Pattern (iii), found only in clade I (*A. reticulata*, *A. serpentaria*), showed *ndhF* extending into IRb—this configuration is novel for Aristolochiaceae. Due to incomplete scaffolds, the junction pattern for *A. californica* (clade IV) remains unresolved.

The plastid phylogeny (Figure 2c; Supplementary Figure S3), inferred from concatenated coding sequences (CDS, tRNA, rRNA, and ORFs), broadly mirrors the nuclear topology. *Thottea* is recovered as sister to *Aristolochia* s.l., with subg. *Siphisia* forming a strongly supported monophyletic clade sister to subg. *Aristolochia*. The major backbone clades (clades I–VII) are largely congruent with those in both nuclear trees. A notable cytonuclear conflict is observed in clade III, where *A. manshuriensis* is recovered as sister to *A. tomentosa*, rather than to *A. macrophylla*. Among Asian clades (V–VII), topological incongruence is more prevalent, with 9 of 53 internal nodes (17%) showing bootstrap support <100 (Supplementary Figure S3). Branch lengths within *Siphisia* remain short, indicating limited plastid divergence (Supplementary Figure S3). GC content analysis across the complete plastomes (Figure 2c) revealed minimal variation among taxa—both in overall genome composition and within specific genomic regions (LSC, SSC, IR), gene categories (CDS), and third codon positions (GC3). This high level of sequence conservation corresponds with the short branch lengths observed in the plastid tree and aligns with patterns seen in the nuclear phylogeny (Supplementary Figure S3), particularly among Asian endemic clades. Notably, A*. griffithii*, which is consistently placed within clade V in the nuclear trees, is recovered outside of this clade in the plastid phylogeny, thereby disrupting its monophyly. To accommodate this topological incongruence, we designate the remaining taxa—still forming a monophyletic group in the plastid tree but now sister to clade VII—as clade Va, and *A. griffithii* as clade Vb (Supplementary Figure S3), which forms a single bifurcating branch sister to clades Va, VI, and VII. This delineation is adopted for clarity in subsequent analyses and discussion.

### 3.3. Cytonuclear conflict and gene–species tree discordance: ILS and introgression

#### 3.3.1. Gene-tree and species-tree discordance results and cause

To investigate sources of topological discordance, we quantified gene–species tree incongruence by comparing each node in the ASTRAL species tree to its corresponding topologies across single-gene trees. Results are shown in Figure 3a, where pie charts at each node represent the proportion of gene trees supporting: the species topology (blue), the dominant conflicting topology (green), the remaining non-dominant conflicting topologies (red), and uninformative gene trees (gray). Despite widespread gene-tree discordance throughout the tree, most major nodes delineating the seven clades established in the nuclear phylogeny showed >50% support from gene trees (blue). One notable exception is node γ (Figure 3a), where non-dominant conflicting topologies (red) outweigh both the species topology and the dominant alternative. This node corresponds to the divergence leading to the three Asian endemic clades (V–VII), which also exhibit coalescent–concatenation incongruence (Supplementary Figure S2) and marked cytonuclear discordance (Figure 2, Supplementary Figure S3).

**Figure 3.**
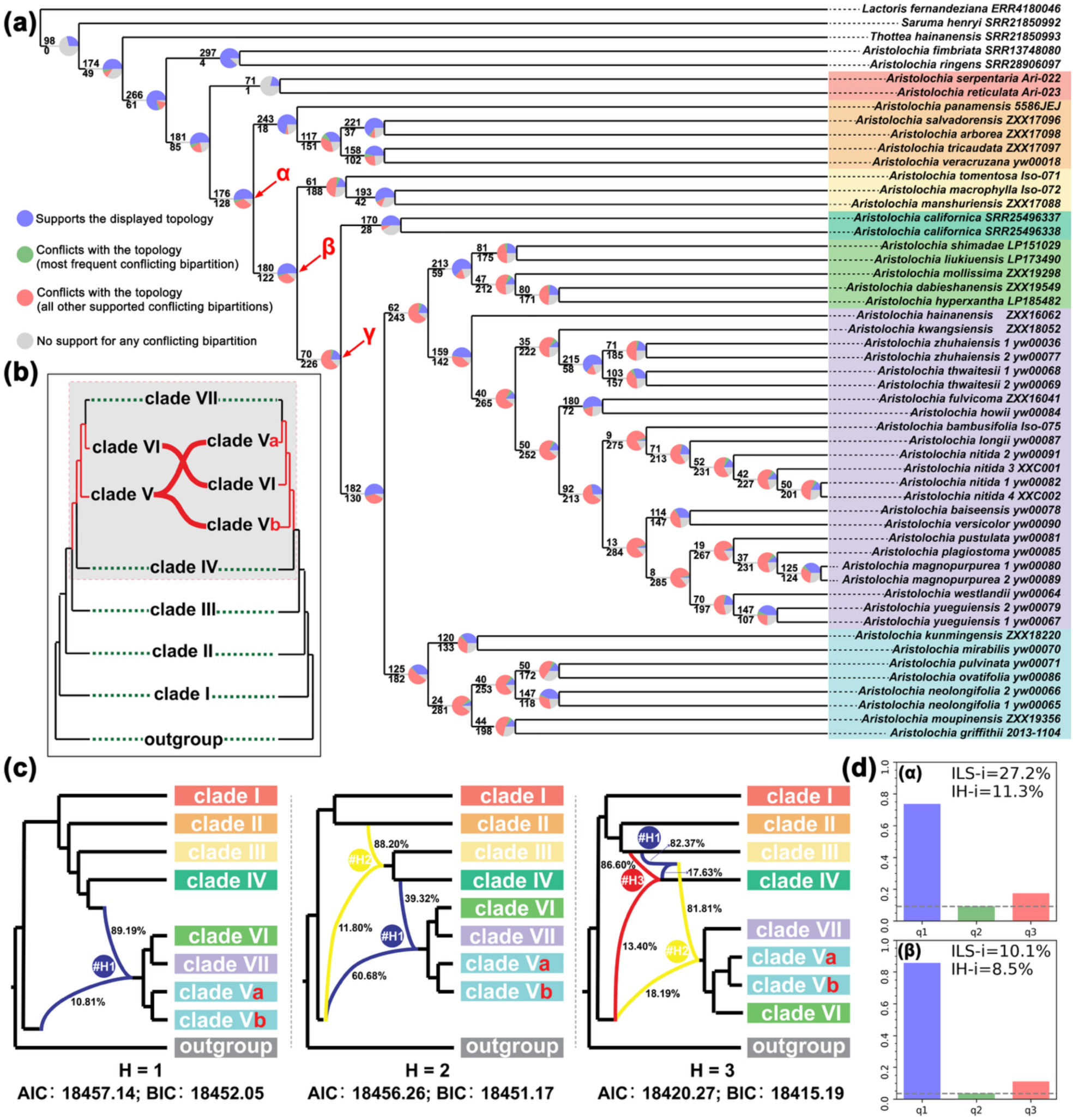
Phylogenomic conflict and reticulation analyses in *Aristolochia* subg. *Siphisia*. **(a)** Gene–species tree concordance analysis. Each node’s topology in the coalescent species tree is compared with individual gene trees. Pie charts indicate the proportion of gene trees that are concordant (q1) or discordant (q2, q3) with the species tree; supporting gene tree counts are shown above (concordant) and below (most frequent discordant) each node. Notably, node **γ**— representing the endemic Asian *Siphisia* clade comprising Clades V, VI, and VII—shows a high degree of conflict with rare dominant alternative topology, suggesting substantial incomplete lineage sorting (ILS). Clade color codes follow the scheme used throughout the main text. **(b)** Cytonuclear discordance between nuclear (coalescent-based, left) and plastome (right) phylogenies. Clade V is consistently monophyletic in both concatenated and coalescent nuclear trees (Figure 1; Supplementary Figure S1); however, *Aristolochia griffithii* (i.e., Clade Vb) is not recovered as monophyletic in the plastome tree (Figure 2c; Supplementary Figure S3). Additional topological conflicts are observed among the endemic Asian clades V, VI, and VII. **(c)** Phylogenetic network inference using PhyloNet (maximum pseudo-likelihood), allowing one to three hybridization events. Curved edges represent inferred hybridization, with associated inheritance probabilities. All models infer reticulation at basal nodes in *Siphisia*, particularly involving North American lineages. No hybridization is detected among endemic Asian clades (V–VII). **(d)** ILS and introgression detection using Phytop. Quartet frequencies (q1: concordant; q2/q3: alternative topologies) are shown along the x-axis for nodes **α** and **β**—the only nodes that yielded signals of hybridization among all tested nodes (Supplementary File S6), corroborating the reticulation events inferred in **(c)**. ILS-i: estimated ILS index; IH-i: estimated introgression index.

In general, support for the species tree declines in the nodes approaching the tips, where discordance is increasingly dominated by a diffuse set of non-dominant conflicting topologies. This pattern reflects reduced phylogenetic signal and elevated gene tree heterogeneity in recent divergences. This pattern, coupled with low node support in phylogenies, strongly suggests incomplete lineage sorting (ILS) as a likely major driver of discordance at shallow nodes.

### 3.3.2. Cytonuclear conflicts and reticulation events

To assess the robustness of the seven major clades delineated above and to explore the causes of cytonuclear discordance observed in the phylogenies, we reconstructed a reduced backbone tree (Figure 3b) representing one species per clade. This comparison confirmed that topological conflicts between nuclear and plastid phylogenies are concentrated within the three most species-rich Asian endemic clades (clades V–VII). In the plastid tree, clade V (here subdivided into Va and Vb, where Va includes clade V taxa as defined by the nuclear phylogeny excluding *A. griffithii*, and Vb comprises only *A. griffithii*) does not mirror the relationships inferred from the nuclear topology. Specifically, Va is recovered as sister to clade VII, and this Va plus VII lineage is sister to clade VI; the combined lineage is then sister to Vb (*A. griffithii*). The remainder of the backbone relationships are congruent across datasets.

To further investigate potential causes of these conflicts, we applied PhyloNet to infer reticulation events (Figure 3c). Under models allowing for 1, 2, and 3 hybridization events, all scenarios recovered ancient introgression events predating the divergence of the Asian endemic clades (V–VII). The model with three reticulation events yielded the best fit (lowest AIC and BIC; Figure 3c, Supplementary File S5), suggesting deep hybridization involving American lineages that may have contributed to the ancestral formation of these Asian clades. However, no reticulation events were inferred within the Asian lineages themselves.

We further quantified the contributions of incomplete lineage sorting (ILS) versus introgression using Phytop. Surprisingly, only two internal nodes—denoted α and β in Figure 3A—showed detectable contributions from hybridization/introgression (13.1% and 8.5%, respectively). These correspond to the basal divergences we inferred within subgenus *Siphisia*, reinforcing the PhyloNet results and suggesting that hybridization occurred early in the lineage’s history. All remaining discordant nodes, including those within clades V–VII, were attributed solely to ILS. This pattern suggests that *Siphisia* underwent rapid radiation, which likely facilitated frequent incomplete lineage sorting while concurrently limiting introgression. In particular, the complete absence of introgression signals within the three Asian clades, despite extensive discordance, highlights ILS as the primary driver of gene–species tree conflict in this lineage (Figure 3D; Supplementary File S6).

### 3.4. Floral morphology and the Aristolochia versicolor species group

#### 3.4.1. Species identity reassessment of Aristolochia versicolor

During sampling preparation, we identified notable discrepancies in the identity of *Aristolochia versicolor* S.M.Hwang, a taxa within clade VII. This prompted a re-examination of type material, field specimens, and literature-based records. *A. versicolor* was originally described by Hwang (1981) based on plants from Xishuangbanna, Yunnan (holotype: HITBC0037109), with two paratypes from Guangxi and Guangdong (Figure 4b,e; Supplementary Table S3). The holotype bears preserved floral material, with the label noting a yellow-green adaxial calyx limb consistent with the original description. In contrast, the Guangxi paratype noted a dark purple flower but lacked reproductive tissues, and the Guangdong paratype was strictly vegetative.

**Figure 4.**
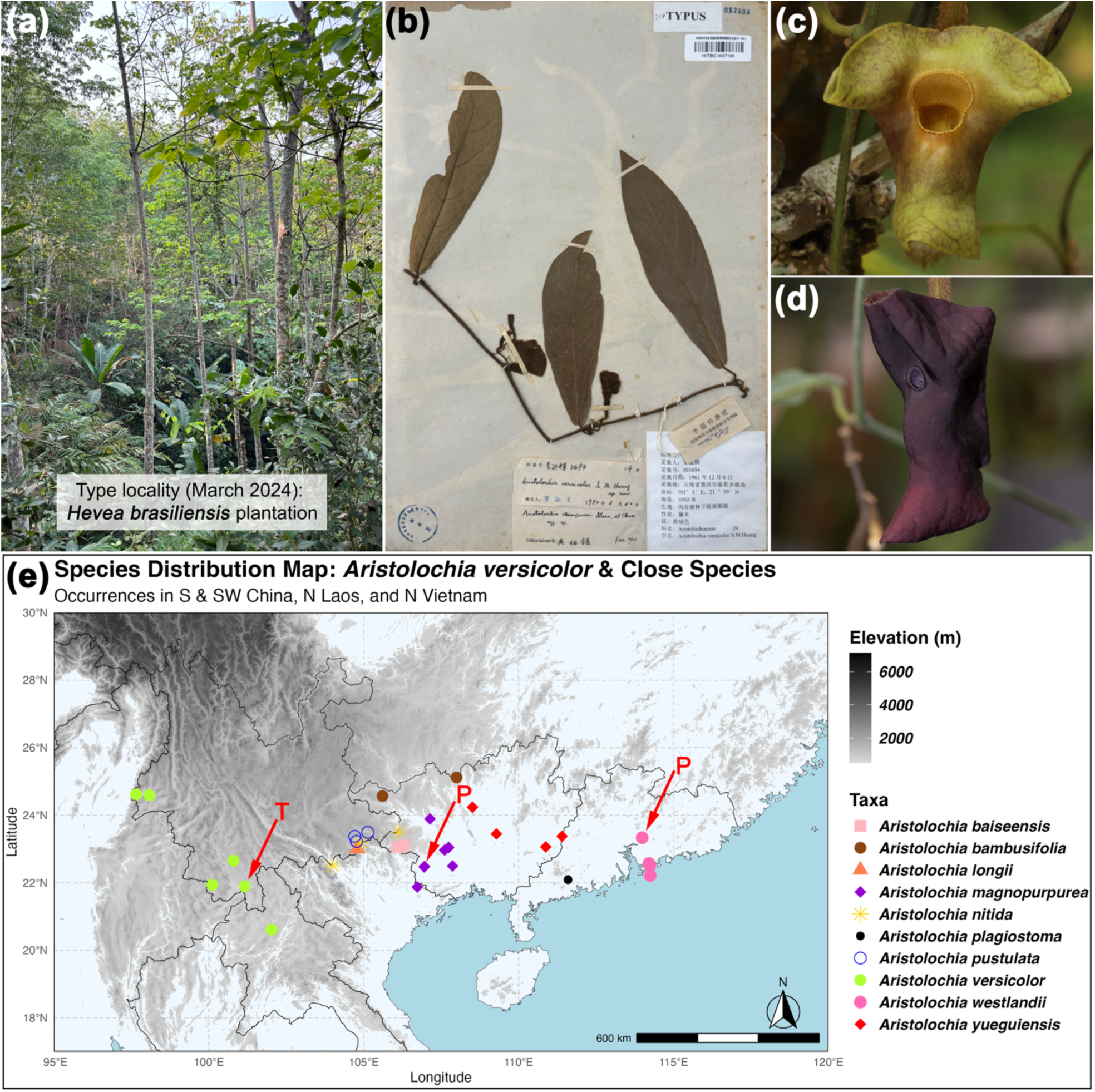
*Aristolochia versicolor* illustration and biogeographic distribution of the *A. versicolor* species group. **(a)** Revisit of the type locality in March 2024, where the original forest has been entirely replaced by artificial plantations. The red arrow indicates rubber trees (*Hevea brasiliensis*, Euphorbiaceae), and the yellow dashed box highlights Tsao-Ko (*Lanxangia tsao-ko*, Zingiberaceae) intercropped in forest gaps. No *Aristolochia* individuals were observed. **(b)** Holotype specimen of *A. versicolor* (voucher barcode: HITBC0037109, XTBG Herbarium). **(c)** Living individuals of *A. versicolor* matching the original description, collected from Pu’er City (coll. No.: *0020200049*) and cultivated in the XTBG Vine Garden, exhibiting floral variation with a purple hue over a green-yellowish base. **(d)** *Aristolochia magnopurpurea* Y.Fan Wang, Z.R.Guo & J.G.Onyenedum, sp. nov., previously misidentified as *A. versicolor* in the literature. It is characterized by a uniformly dark purple adaxial calyx limb and is primarily distributed in the karst limestone regions of western Guangxi Province and northeastern Vietnam. **(e)** Distribution map of the *A. versicolor* species group. Green circles indicate confirmed occurrences of *A. versicolor*, primarily within tropical rainforest climate zones from northern Laos to western Yunnan. “T” marks the type locality; “P” indicates the paratype localities originally described in two geographically isolated provinces—Guangxi (now confirmed as *A. magnopurpurea*) and Guangdong (now confirmed as *A. westlandii*). Other species within the *A. versicolor* species group are color-coded on the map to indicate their respective distributions.

Extensive fieldwork in March 2024 revealed that the original type locality had been fully converted to plantations (Figure 4a), but morphologically consistent populations with yellow-green calyx were identified and sampled from Pu’er and Dehong (Yunnan), and northern Laos (Figure 4e). These populations occur in tropical monsoon forests (Cao et al., 2006) and share climatic and morphological traits with the type.

Literature review revealed additional inconsistencies. Accessions from Vietnam (Do and Hoang, 2022) and Laos (Le and Do, 2022) attributed to *A. versicolor* exhibit uniformly dark purple calyces and small throat openings (Figure 4d), inconsistent with the yellow-green morphotype. In Le and Do (2022)’s case, a voucher label indicated a greenish-yellow flower, but was published alongside a photograph of a purple-flowered plant, raising concerns of a chimeric presentation. In contrast, Liao et al. (2021) and Zhu and Ma (2022) documented yellow-green floral morphotypes with larger throats, consistent with the type description. Across all observed specimens, one trait remained consistent: the reflexed calyx limb during anthesis (Figure 4c–d), which was also present in *A. westlandii* (Figure 1q) and five undescribed taxa collected during this study (Figure S4), suggesting potential shared ancestry.

Phylogenomic analyses confirmed the observed morphological and geographic patterns. In both the coalescent-based (Figure 1) and concatenated supermatrix (Figure S1) nuclear phylogenies, *Aristolochia versicolor* was recovered within a strongly supported monophyletic clade comprising 15 accessions representing 10 taxa, including two species with non-reflexed calyces: *Aristolochia bambusifolia* C.F.Liang ex H.Q.Wen and *Aristolochia plagiostoma* (*Isotrema plagiostomum* X.X.Zhou & R.J.Wang) (Figure 1; Supplementary Figure S1; Supplementary Figure S4). This entire lineage received full node support (LPP = 1.0; BS = 100). Within the clade, *A. versicolor* was consistently resolved as sister to *A. baiseensis* (in prep.), a morphologically distinct but undescribed species (Supplementary Figure S4). This placement renders the remaining taxa in the clade paraphyletic with respect to *A. versicolor*. In the plastid phylogeny (Figure 2; Supplementary Figure S3), *A. versicolor* and *Aristolochia baiseensis* Su Y.Nong, H.C.Xi & Y.S.Huang, sp. nov. (manuscript in prep.) were also recovered as sister taxa with full support (BS = 100), forming a distinct lineage. However, the full set of ten taxa defined in the nuclear phylogeny was not recovered as monophyletic in the plastid tree, indicating a case of cytonuclear discordance.

Together, these results support the recognition of *A. versicolor* as a narrowly distributed taxon confined to tropical monsoon forests of southwestern Yunnan and northern Laos, characterized by yellow-green flowers. Other populations previously identified as *A. versicolor*, particularly those bearing dark purple flowers or exhibiting divergent floral morphologies, represent cryptic new species that warrant formal description. Furthermore, these taxa— together with *A. bambusifolia* and *A. plagiostoma*—constitute a well-supported and morphologically coherent clade, herein defined as the *Aristolochia versicolor* species group.

#### 3.4.2. Calyx tube epidermis examination

We examined all *Aristolochia* subg. *Siphisia* taxa included in this study using flowers from living collections, which allowed not only careful morphological observation but also temporal tracking of reproductive development during anthesis. Strikingly, without exception, all sampled taxa: from the morphologically distinct herbaceous clade I to the highly diverse clade VII, displayed a glabrous upper calyx tube. This region, homologous to the upper tube of subg. *Aristolochia* and subg. *Pararistolochia*, consistently lacks the downward-pointing trichomes known to be involved in pollinator trapping in those subgenera. Instead, in *Siphisia*, this surface is entirely smooth, suggesting a different perianth–pollinator interaction mechanism. Representative images are shown in Figure 5 (panel a–d). In particular, *A. versicolor* (Figure 5a) and its pressed voucher’s adaxial epidermis under 20× magnification (Figure 5b) show papillate cell patterns but no trichome development. The same glabrous condition was observed in all *Siphisia* species included in this study: for example, *A. bambusifolia* (Figure 5c) and *A. plagiostoma* (Figure 5d), confirming that the absence of this key trapping trait is widespread across the subgenus and may represent a defining synapomorphy for *Siphisia* as a whole.

**Figure 5.**
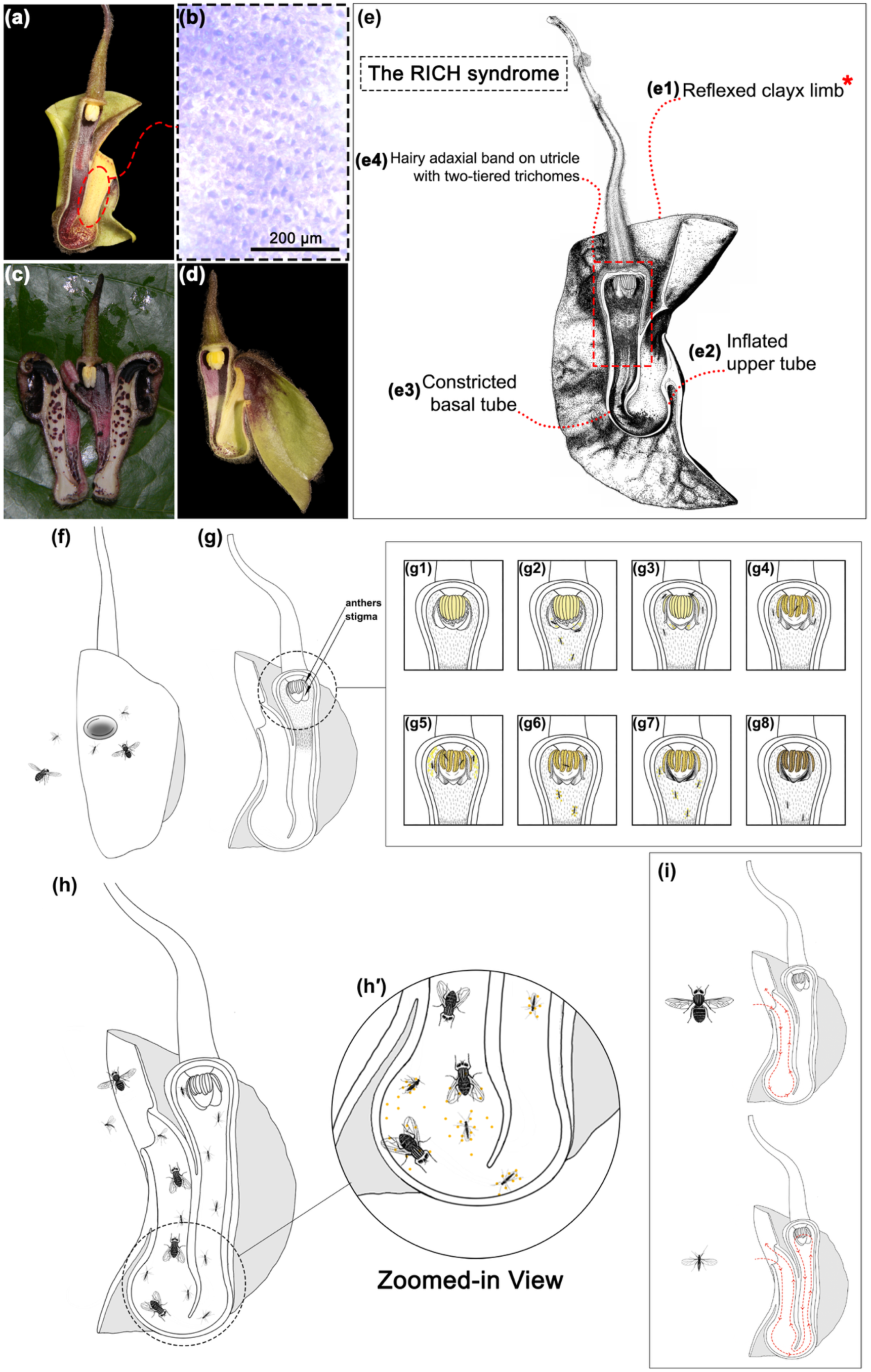
Morphological and functional specialization in *Aristolochia* subg. *Siphisia*, with emphasis on the *A. versicolor* species group. **(a)** Longitudinal dissection of *A. versicolor*. **(b)** Stereomicroscopy of the adaxial surface of the upper tube in *A. versicolor*, showing an absence of trichomes and presence of smooth, glabrous epidermal cells with weakly papillate texture, producing a waxy surface. **(c–d)** *A. bambusifolia* and *A. plagiostoma*, respectively, both lacking a reflexed calyx limb. **(e)** The “RICH” syndrome, a floral character complex unique to the monophyletic *A. versicolor* group (Figure 2), composed of four synapomorphies: **(e1)** a reflexed calyx limb (an incomplete synapomorphy, marked with a red asterisk, absent in *A. bambusifolia* and *A. plagiostoma*); **(e2)** an inflated upper tube; **(e3)** a constricted basal tube entrance hypothesized to filter non-target pollinators; and **(e4)** a hairy adaxial band on the utricle surrounding the gynostemium, composed of two distinct trichome tiers—potentially serving separate functions in nectar secretion (lower tier) and pollen transfer (upper tier). **(f–i)** Schematic representation of the pollination process in the *A. versicolor* species group: **(f)** Pollinator attraction and entry into the floral tube. **(g)** Typical gynostemium structure and developmental chronology in *Siphisia*. **(g1–g4)** Pistillate phase: **(g1)** Stigma turgid and receptive; **(g2)** Pollen-carrying pollinators enter; **(g3)** Stigma begins to contract and dry after pollen reception, while pollinators remain trapped; **(g4)** Anthers begin to separate and protrude. **(g5– g8)** Staminate phase: **(g5)** Pollen is released; **(g6)** Pollinators acquire pollen; **(g7)** Stigma fully contracts and hardens; **(g8)** Pollinators exit carrying pollen. The yellow-colored structures represent the stamens. Only one stigma lobe (of three) and its two laterally attached anther pairs (of six total) are shown in the illustration, progressing from a premature state (light yellow, **g1– g3**) to active pollen dehiscence (bright yellow, **g4–g7**), followed by pollen exhaustion or degeneration **(g8)**. **(h)** Floral morphology modulating insect entry and exit based on body size. **(h′)** Close-up of the geniculation site, showing physical restriction that excludes non-target insects. **(i)** Hypothetical pollination trajectories: upper path—non-target insects trapped in the upper tube; lower path—effective pollinators reaching the gynostemium and exporting pollen. Note: Pollinator illustrations are schematic and hypothetical; interactions are inferred from floral morphology and remain to be verified through field-based pollination studies.

#### 3.4.3. The “RICH syndrome”

To further understand the monophyletic *A. versicolor* species group defined previously, we examined floral characters across its constituent 10 taxa to identify potential lineage-specific synapomorphies. This led to the recognition of the “RICH syndrome,” a unique combination of four floral traits that consistently define this group (Figure 5). These include: (1) a reflexed calyx limb, present in all group members except *A. plagiostoma* and *A. bambusifolia*; (2) a conspicuous inflation at the junction of the upper and basal tubes, although less pronounced in *A. nitida* sp. nov.; (3) a sharply constricted basal tube, forming a narrow aperture contrasting with the inflated upper tube; and (4) a villous adaxial utricle surface, forming two clearly defined bands surrounding the gynostemium. This villous region frequently extends below the gynostemium, a pattern not observed in other *Siphisia* taxa, where the utricle hair bands are typically flat and the two-tiered structure is inconspicuous.

We additionally documented a temporal shift in gynostemium and utricle hair development across *Siphisia*, indicative of protogyny. Figure 5 (panels g1–g8) illustrates this sequential maturation, consistent with a pollen reception–release timing characteristic of protogynous dichogamy, a reproductive strategy also widely reported in other magnoliids (Endress, 2010).

## 4. Discussion

### 4.1. Systematics and Circumscription of Subgenus Siphisia

This study presents the first well-resolved phylogenomic framework for *Aristolochia* subg. *Siphisia*, combining broad taxon sampling with a universal genome-wide nuclear probe plus complete plastomes to achieve robust species-level resolution. Earlier efforts, whether based on a few loci (e.g., *trnL-trnF*, *rbcL*, *phyA*, *ITS2*) or chloroplast genome assemblies, were constrained either by sparse sampling or by insufficient phylogenetic resolution, resulting in poorly resolved topologies or unsupported polytomies (e.g., Bai et al., 2023; Jost and Wanke, 2024; Li, 2019; Ohi-Toma et al., 2006, 2021; Wanke et al., 2006, 2017; Xu et al., 2023; Zhu et al., 2019b). By contrast, our analysis resolves seven well-supported clades within *Siphisia*, filling a major gap in the systematics of this morphologically and ecologically diverse group.

We adopt the coalescent-based species tree as the primary reference topology. Nuclear genome-wide data reflect biparental inheritance and allow modeling of incomplete lineage sorting (ILS), making them theoretically and empirically more suitable for resolving species-level relationships in groups undergoing rapid radiations (Degnan and Rosenberg, 2009). While our concatenated tree yields a highly supported and largely congruent backbone (Figure S1), coalescent inference avoids potential biases arising from gene-tree heterogeneity (Degnan & Rosenberg, 2006; Mirarab et al., 2021). Importantly in our analysis, both coalescent and concatenated trees (Figure 1, Supplementary Figure S1, Supplementary Figure S2) yielded broadly congruent topologies, yet we observed minor but meaningful conflicts in the deeper nodes of Asian endemic clades—precisely where gene-tree discordance and phylogenetic conflict are inferred (Figure 4a).

Li (2019) provided one of the most comprehensive early frameworks for *Siphisia*, sampling 80 accessions across 62 taxa using five plastid and three nuclear loci. Her study offered valuable insights into clade structure and biogeographic trends, laying important groundwork for later efforts. However, relationships among the major Asian endemic lineages remained unclear. The group corresponding to our clades V–VII was recovered as a single polytomy, with clades VI and VII forming an unresolved cluster placed as sister to clade V. As Li also noted, the disparity in species richness: roughly 80% of *Siphisia* diversity occurring in Asia, may reflect deeper evolutionary asymmetries. Our genome-wide results clarify this region of the tree with strong support: clade VII, distributed primarily in southwestern China and Indochina, is consistently recovered as sister to the Sino-Japanese clade VI, rather than to clade V, which centers in SW China and the Hengduan region.

We further refine the circumscription of these seven clades by integrating geographic distribution and morphological synapomorphies (Figure 1). Following Wu and Wu (1996), the clades align with distinct floristic regions: (1) eastern North America (clade I); (2) Central America and southern Mexico (clade II); (3) a classic eastern Asia–eastern North America disjunct lineage spanning northeastern Asia and the southeastern U.S. (clade III); (4) California (clade IV); (5) the tropical montane zones of the Indo-Burma biodiversity hotspot, including the southern Himalayas and Hengduan region (clade V); (6) the Sino-Japanese subtropical region (clade VI); and (7) tropical southern China south of 22°N, extending into Indochina and Malaysia (clade VII). These disjunctions likely reflect a combination of ancient intercontinental dispersals (e.g., via the Bering land bridge) and region-specific radiations shaped by tectonic uplift and climatic oscillations throughout the long-term evolution (Tiffney & Manchester, 2001; Wen et al., 2016).

Notably, several clades exhibit diagnostic root morphotypes—traits largely overlooked in prior treatments of *Aristolochia*. For example, clade VII taxa typically inhabit limestone karst and develop fusiform or moniliform tuberous roots that anchor plants into rocky substrates. In contrast, clade VI taxa, found in loamy, clay-rich soils of the Sino-Japanese region, possess creeping rhizomes. These morphologies may reflect adaptive responses to edaphic conditions, aligning with the role of geophytism in diversification and persistence under selective pressure (Tribble et al., 2021). While floral characters are typically emphasized in *Aristolochia* taxonomy, here, we find those traits to play a secondary role in our systematic framework due to their ecological plasticity. But this does not mean floral traits are not informative. In fact, it is the presence of the RICH syndrome floral synapomorphies defines the *A. versicolor* species group (see Section 3.4). Future studies aiming to expand taxon sampling should prioritize living material to capture floral dynamics, particularly in traits like anthesis timing and trichome development that are difficult to observe in dried specimens.

It is also worth noting that plastome-based phylogenies, even those based on complete chloroplast genomes, show limited resolution for species-level relationships within *Siphisia*, particularly among the Asian endemic clades where cytonuclear discordance is pronounced (Figure 2, Figure 3, Supplementary Figure S3). While plastid genomes offer valuable insight into matrilineal inheritance patterns (Korpelainen, 2004), they are best treated as complementary sources of information, rather than as stand-alone frameworks for primary phylogenetic inference. In our study, the plastid tree failed to recover several species groups that were consistently supported by nuclear data and easily diagnosable by morphology (Figure 2), despite full plastome recovery and synteny validation. This highlights the need to interpret organellar phylogenies with caution and emphasizes the distinct evolutionary histories, and potential conflicts, between nuclear and organellar genomes.

Looking forward, the systematics of *Siphisia* could be further refined through clade-specific probe design and higher-resolution sampling at the population level. The recent availability of a chromosome-level *Siphisia* genome of *A. manshuriensis* (Hu et al., 2025), opens new possibilities for custom single-copy nuclear markers (Chamala et al., 2015), improved SNP calling, and population genomics. As our study provides nearly half of the described *Siphisia* species with phylogenomic data, it lays a strong foundation for such advances.

### 4.2 Evolutionary Dynamics in Siphisia: Speciation and Hybridization

Li (2019) proposed that *Siphisia* species exchanges among the Malaysia, Sino-Japanese, and Sino-Himalayan floristic regions were frequent during the late Oligocene warming and the Miocene Climatic Optimum, driven by a combination of climatic and geological events (Zachos et al., 2001). These diffusion-based processes were suggested to underlie its remarkable species richness of in Asia. However, our assessments of gene flow and lineage divergence challenge this view. Specifically, our analyses show that hybridization is unlikely to have played a major role in the diversification of *Siphisia*. Nearly all hybridization signals are confined to non-Asian lineages, mostly among Central and North American taxa, and even these are limited and weak (Figure 3, Supplementary File S6). Phylogenetic network analyses suggest that these signals may reflect historical admixture between ancestors of currently disjunct American species, rather than recent or ongoing hybridization (Figure 3).

The striking contrast between the near-complete absence of introgression in the speciose Asian clades and their exceptionally high species richness prompts deeper inquiry into alternative drivers of diversification. Notably, incomplete lineage sorting (ILS) was pervasively detected across our nuclear dataset, accompanied by extremely short internal branch lengths in both nuclear and plastid phylogenies. Together, these patterns are indicative of rapid cladogenesis, with multiple speciation events occurring within a narrow temporal window. A possible contributing factor, as proposed by Watanabe et al. (2008) based on the *A. kaempferi* complex (corresponding to our clade VI), is strong geographic structuring of genetic diversity driven by Pleistocene climatic oscillations. Cycles of range expansion and contraction during glacial–interglacial periods likely promoted population isolation and divergence, facilitating lineage differentiation and the accumulation of biodiversity.

Two key points arising from these patterns warrant further investigation. First, what mechanisms are responsible for limiting hybridization, especially given that *Aristolochia* s.l. is known to hybridize naturally and is easily crossed in cultivation (Blanco, 2005). Notably, most confirmed hybrid cultivars involve species of subg. *Aristolochia*, with few, if any, cases documented in *Siphisia*. This discrepancy may reflect intrinsic genomic barriers. Cytological studies confirmed that *Siphisia* underwent whole-genome duplication (WGD) (Hu et al., 2025), with many species exhibiting doubled chromosome numbers 2n = 28/32 (Berjano et al., 2009; Fedrov, 1974; Gregory, 1956; Ohi-Toma et al., 2006; Sugawara et al., 2001) compare to subg. *Aristolochia* (2n = 6/12/14/16) (Li, 2019). Such duplication may contribute to reproductive isolation via postzygotic barriers or meiotic incompatibilities, effectively suppressing introgression even among sympatric taxa. Supporting this hypothesis, the study of the *A. kaempferi* complex by Watanabe et al. (2008) detected asymmetric prezygotic isolation, despite the close geographic proximity of the taxa involved.

Second, if extensive ILS is present, as our low-copy nuclear gene trees suggest, how do *Siphisia* species maintain such clearly divergent and diagnosable morphologies, particularly in perianth structure? The striking floral variation across *Siphisia*, which forms the basis of most taxonomic diagnoses, appears decoupled from the underlying stochasticity of gene trees. This raises the possibility that a small number of regulatory loci or structural genes may control perianth development, potentially under strong selection. Developmental shifts or ecological specialization in pollination systems may further accelerate divergence in these traits, facilitating speciation despite a background of shared ancestral polymorphism.

While our current dataset offers only preliminary insight into these dynamics, it establishes a foundation for future research. Population-level genomic sampling will be essential to more precisely quantify gene flow, test for cryptic hybridization, and resolve the timing and mechanisms of lineage divergence. In parallel, comparative studies on reproductive genetics and cytogenetics, particularly those examining meiosis, pollen viability, and embryo development across putative barriers may elucidate the genomic architecture underlying *Siphisia*’s remarkable diversification.

### 4.3. Pollination Biology in Siphisia: Toward a Novel Mechanistic Framework

The near-complete absence of introgression and hybridization signals in our nuclear dataset—despite frequent sympatry among *Siphisa* species (Zhang et al., 2024)—raises an important question: what reproductive strategies are in place that limit interspecific gene flow? On a broader scale, this points toward structural or ecological mechanisms that enforce reproductive isolation, and we argue that the answer lies in both conserved reproductive traits and lineage-specific adaptations.

Across *Aristolochia*, some general mechanisms are shared across many lineages in Magnoliids (Bertin and Newman, 1993; Endress, 2010). Protogyny (Figure 5), for example, is widespread and enhances outcrossing by temporally separating female and male phases, thus minimizing selfing (Alpuente et al., 2023; Matallana-Puerto et al., 2024). While this mechanism likely underpins many *Aristolochia* species, species-and clade-specific floral morphologies and behaviors also likely play a critical role in mediating pollinator specificity and pollen flow. Notably, nearly all detailed pollination studies within *Aristolochia* have focused on subg. *Aristolochia* (e.g., Alpuente et al., 2023; Berjano et al., 2009; Blatrix et al., 2024; Burgess et al., 2004; Erbar et al., 2017; Oelschlägel et al., 2009, 2015; Park and Kim, 2023; Rulik et al., 2008; Rupp et al., 2021; Trujillo and Sérsic, 2006). These studies have revealed a wide array of highly specialized strategies: from brood-site mimicry (Rupp et al., 2021) to direct insect competition (Berjano et al., 2009) between sympatric species. Yet, the underlying floral architecture is consistent across these systems, centered on a well-known trap-and-release mechanism (Oelschlägel et al., 2009). In *Aristolochia* s.s., the typical pollination mechanism involves a utricle at the base, which houses the gynostemium. The orientation of the stigma can be highly variable: skewed downward, horizontal, or upward, depending on the species and taxonomic section. Above the utricle is a narrow, elongated tube lined with downward-pointing trichomes (González and Stevenson, 2000b). These trichomes serve as the key structural element enabling successful entrapment: as many pollinators exhibit positive phototaxis or upward movement tendencies (Jander, 1963; Jékely, 2009), the trichomes prevent their escape, retaining them inside the floral chamber until the pollination cycle is complete.

Surprisingly, *Siphisia* has received comparatively little attention, despite its expanding diversity. González and Pabón-Mora (2015) noted in a commentary that the floral mechanism in *Siphisia* likely diverges from that of *Aristolochia* s.s., suggesting the possibility of an alternative trapping and insect-release strategy. Erbar et al. (2017) examined three Central American fungus-imitating *Siphisia* species (corresponding to our clade II) and reported notable differences in nectary structure and secretion patterns, though they did not investigate the entrapment mechanism. Nakonechnaya et al. (2020, 2008) studied floral morphology in *A. tomentos*a and *A. manshuriensis*, but again, the pollination biology remained largely inferential. Overall, taxonomic and biogeographic bias in the literature has left most of *Siphisia* underexplored.

Our own morphological survey of *Siphisia* reveals a structural basis for this knowledge gap: the trichomes central to entrapment in *Aristolochia* s.s. are absent from the upper calyx tube across all *Siphisia* species examined (Figure 5i-iv). Instead, these surfaces are glabrous and either waxy or velvety in texture. Additionally, while *Aristolochia* s.s. flowers possess a single continuous tube, *Siphisia* flowers have a bipartite U-shaped tube with a basal and upper segment, meeting at a geniculation point (González and Stevenson, 2000b; Zhu et al., 2019b). The gynostemium in *Siphisia* is also superior and pendulous, positioned at the distal terminus of the floral tube—a placement that requires pollinators to navigate a longer and more complex path to reach the reproductive organs. This floral design suggests that *Siphisia* may have evolved a fundamentally different pollination strategy: one not reliant on physical entrapment but on structural navigation or reward-driven retention (e.g., nectar guides). The highly elongated, curved perianth tube alone may hinder rapid insect movement, enough serving as a passive trap. This hypothesis aligns with the fly (Diptera)-centric pollination observed in previous studies (Correns, 1891, 1892; Faegri and van der Pijl, 1979; Hildebrand, 1867; Proctor et al., 1996). Dipterans may be well-suited to navigate these long, narrow tubes, especially if visual and olfactory cues are also effectively employed.

In the *A. versicolor* species group, we propose a more refined mechanism: a pollinator filtering system driven by the four synapomorphies of the RICH syndrome (Section 3.4.3). Among these, the most functionally significant appears to be the constriction at the basal tube (Figure 5). We posit that pollinators first accumulate in the inflated upper tube but must squeeze through a narrow passage to reach the utricle, where reproductive organs and possible nectar rewards reside. This narrowing likely acts as a filter, excluding insects above a certain size threshold and optimizing pollen transfer by targeting specific functional pollinators (Figure 5). This design also could balance attraction and selectivity. By keeping the floral entrance (i.e., calyx throat) open, volatile dispersal and visual cues remain unhindered, while the downstream constriction enforces size-based selection to exclude unwelcome intruders. This dual-stage design may enhance pollination efficiency while minimizing the energetic cost of indiscriminate visitor access.

Of the four RICH traits, calyx reflexion is the most conspicuous morphologically but may play a less central role in the process. Two taxa within the group lack this trait but retain the other three synapomorphies (Supplementary Figure S4), suggesting possible convergence or trait loss. Moreover the two-tiered hairy band surrounding the gynostemium likely has functional significance. The lower band, often waxy and wet (Supplementary Figure S4), may serve as a nectary zone, while the upper band, composed of appressed trichomes, likely facilitates pollen adhesion through increased friction.

These floral innovations may reinforce reproductive isolation. Our gene-tree discordance analysis indicates that nearly all phylogenetic incongruence stems from incomplete lineage sorting rather than hybridization (Figure 3). The absence of introgression signals supports the hypothesis that historical gene flow has been limited or absent, potentially due to pollinator filtering strategies in *Siphisia* that restrict genetic exchange—even among sympatric taxa with overlapping flowering periods. By targeting narrow pollinator sets, these species likely maintain reproductive exclusivity and promote lineage divergence.

While these hypotheses are consistent with floral structure and phylogenetic patterns, field-based validation remains essential. Pollinator capture, behavioral studies, and controlled pollination experiments will be critical to fully elucidate these mechanisms. Alternatively, the RICH syndrome may not represent adaptive innovation per se, but rather a constrained evolutionary workaround to ecological deterioration. As Jacob (1977) famously noted, evolution often works not as an engineer but as a tinkerer—repurposing existing materials to produce functional, though sometimes suboptimal, solutions. The enclosed, connate calyx—or perianth—seen across *Aristolochia* s.l. may have initially co-evolved with a now-diminished pollinator guild, dragging down with it the floral architectures that had become dependent on such interactions. Given the irreversible nature of floral fusion, neither structural reversal nor the recruitment of new pollinators would have been easily attainable. Thus, piecemeal modifications emerged as compensations—structural improvisations to enforce arthropod climbing behavior, increase tactile contact, and maximize pollen deposition and uptake during increasingly infrequent pollinator visits. In this view, natural selection did not reengineer but favored partial refinements just sufficient to maintain reproductive function under collapsing mutualisms.

Indeed, elaborations like trichome bands and narrowed tubes can fit into the classic explanation of exaptations: features originally evolved for unrelated purposes but secondarily coopted to maintain function under shifting ecological contexts (Gould and Vrba, 1982). Traits now facilitating pollen adhesion or controlled insect movement might have had otherwise-intended developmental or structural origins in their ancestral state. The five newly described taxa (detailed in the next section of this study), bearing such morphologies, are also facing alarming conservation statuses, hinting that what we observe may not be innovation, but a form of architectural endurance. Just as genetic variance governs adaptive potential (Ohno, 1970), the phenotypic plasticity of floral traits can constrain or enable evolutionary paths (Gould and Vrba, 1982). *A. versicolor* species group may be nearing the edge of that flexibility, where modifications become tactical responses rather than forward leaps. In either case, *Siphisia* has clearly diverged from the canonical trap-and-release pollination model of *Aristolochia* s.s., developing clade-specific, structurally distinct systems molded by ecological niche and historically accumulated genetic differences. To fully decipher how these flowers function, ecological research must return to the field—on the ground, where plants and environments act as an integrated whole.

## 5. Taxonomic treatment

5.1. *Key to Aristolochia versicolor and closely related species*

1a. Calyx limb not reflexed 2

1b. Calyx limb reflexed backward, partially or completely enclosing the perianth tube from behind 3

2a. Calyx limb strongly reduced and involute, forming a minute, inward-curled structure……….. Aristolochia bambusifolia

2b. Calyx limb connate into a slightly oblique, cylindrical tube, with the distal end appearing truncate ***A. plagiostoma***

3a. Leaf blade linear to lanceolate 4

3b. Leaf blade elliptic or broadly ovate 5

4a. Calyx limb (adaxial surface) white with conspicuous reticulate venation, reflexed then coiled forward to form an alate structure ***A. westlandii***

4b. Calyx limb (adaxial surface) dark purple, lacking visible venation, reflexed backward and closely appressed to the tube, without a secondary forward coil ***A. yueguiensis***

5a. Limb surface immediately below the throat densely covered with papillate or pustulate projections; throat margin obscured by folded or undulate calyx limb ***A. pustulata***

5b. Throat margin clearly visible, not obscured by calyx folds 6

6a. Calyx limb adaxially whitish with dense purple spotting ***A. baiseensis***

6b. Calyx limb adaxially uniformly colored or mottled, lacking distinct dotted pattern 7

7a. Calyx limb adaxially extremely waxy and lustrous; throat broadly elliptic; calyx tube adaxially dark purple near the throat, but bright yellow throughout the rest ***A. nitida***

7b. Calyx limb adaxially velvety in texture; throat round or terete in outline 8

8a. Calyx limb adaxially greenish-yellow as the base color, sometimes with a purplish hue; throat broad, pure yellow or yellow with sparse, minute orange flecks ***A. versicolor***

8b. Calyx limb adaxially brownish red or purple 9

9a. Calyx limb large (4.5–7 × 5–7 cm), adaxially dark purple, becoming darker toward the throat A. magnopurpurea

9b. Perianth very small (1.6–2.7 × 1.6–2.7 cm), brownish red with sparse yellowish flecks; throat extremely small (3.0–5.4 mm) ***A. longii***

### 5.2. Emended Circumscription of Aristolochia versicolor

***Aristolochia versicolor*** S.M.Hwang, 1981 **Figure 6**

**Figure 6.**
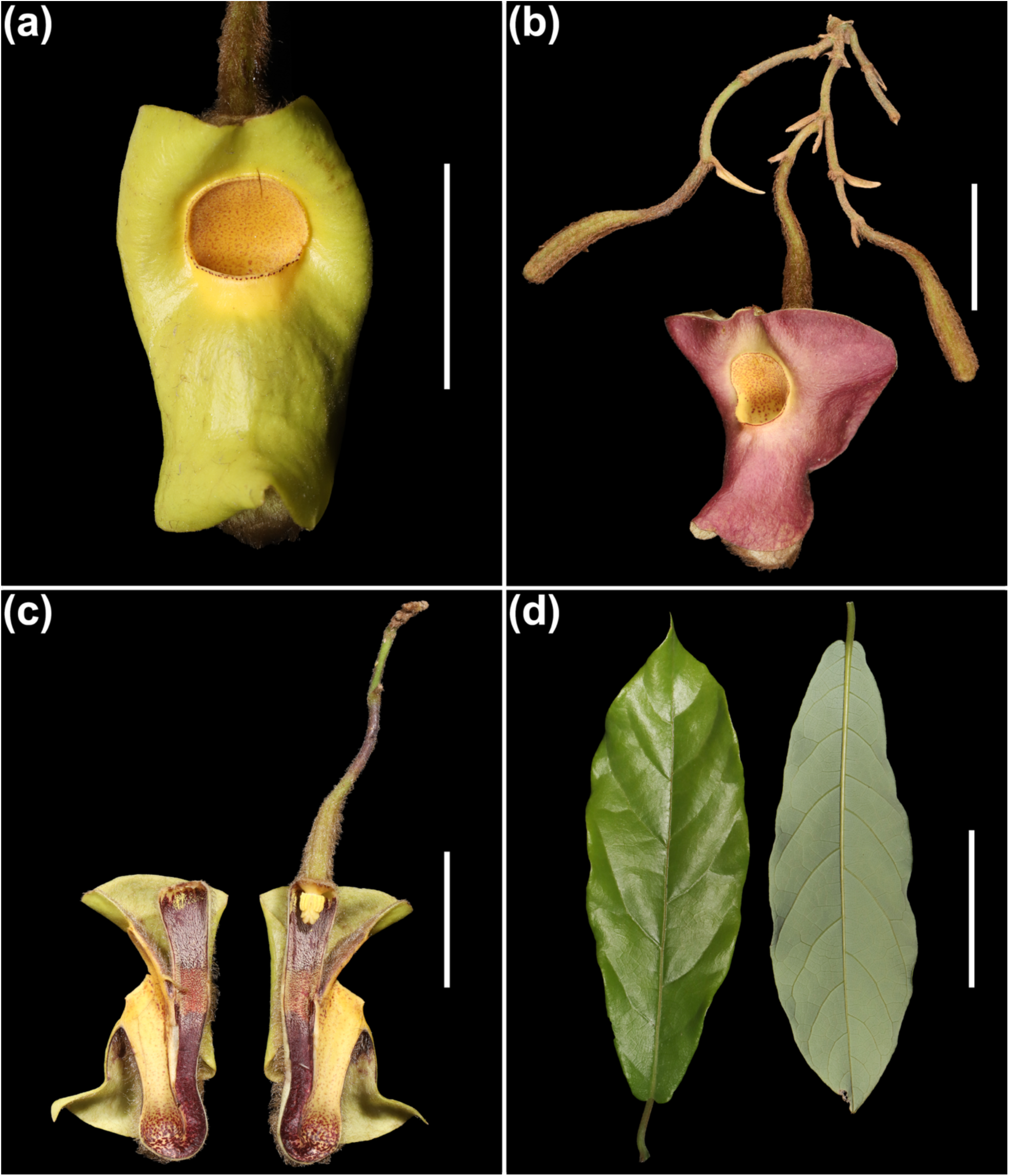
*Aristolochia versicolor* S.M. Hwang, illustration. **(a)** Habit, showing typical greenish-yellow calyx limb. **(b)** Variation in limb coloration with purplish hue occasionally observed; this variation is not developmental. **(c)** Longitudinal dissection of the flower, showing adaxial structure of the calyx tube. **(d)** Adaxial and abaxial surfaces of the leaf. Scale bars: (**a–c)** = 2 cm; **(d)** = 5 cm.

**Emended description:—** Liana. Older stems glabrous, covered with thickened brown bark, irregularly and longitudinally fissured. Leaves coriaceous or papery, narrowly oblong-lanceolate; apex acute; base narrowly cordate or narrowly auriculate; adaxial surface glabrous; abaxial surface glaucous, pilose along the veins, becoming glabrescent with age. Lateral veins 4– 6 on each side, curving near the margin and forming anastomoses. Petioles 1–2 cm long, finely hairy, swollen at the base, slightly twisted and stout. Inflorescences axillary, cymose, 1–6-flowered; pedicels 2–3 cm long, rusty-villous, twisted, each flower 1-bracteate; bracteoles sessile. Calyx limb adaxially 3.0–3.5 × 3.5–4.0 cm, bright yellowish green, with some individuals exhibiting a purplish hue overlaying the yellow-green base color; surface waxy or velvety, glabrous. Upper tube 2.5–3.0 mm long; basal tube 3.0–3.8 mm long; limb reflexed, 3-dentate. Calyx mouth orbicular, 0.8–1.2 mm in diameter. Ovary 6-locular. Stamens 6, with oblong anthers arranged in pairs, grouped in 3 sets opposite the lobes of the column; column apex trilobed. Capsule elliptic, 5–8 cm long, 1.5–2 cm wide, yellow-green, hexagonal, turning dark brown at maturity. Seeds ovate, brown.

**Amended distribution and habitat:—** *Aristolochia versicolor* is currently known only from the tropical rainforest monsoon zone of northern Laos and southwestern Yunnan, China, particularly in regions such as Pu’er, Xishuangbanna and Dehong (Figure 4e). Although previously reported from Guangxi, Guangdong, and Northen Vietnam, these records have been re-evaluated and are now attributed to distinct, undescribed species. No verified occurrences of true *A. versicolor* have been confirmed outside the aforementioned areas.

### 5.3. Descriptions of new species

*Aristolochia magnopurpurea* Y.Fan Wang, Z.R.Guo & J.G.Onyenedum, **sp. nov. Figure 7**.

**Figure 7.**
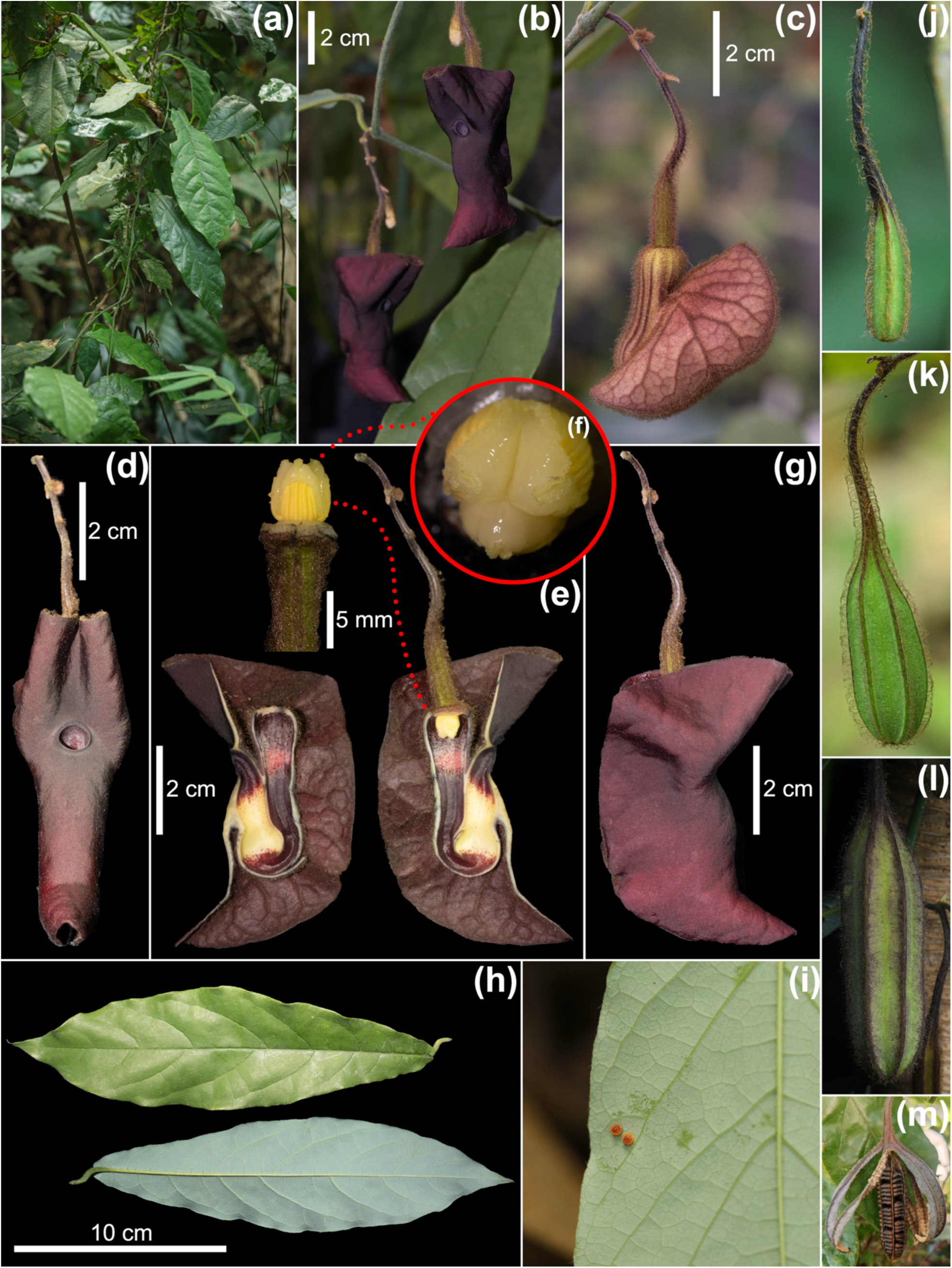
Illustration of *Aristolochia magnopurpurea* Y.Fan Wang, Z.R.Guo & J.G.Onyenedum, sp. nov. **(a)** Plant in situ. **(b)** Axillary racemose inflorescence; flowering initiates from the terminal flower, followed by sequential development of buds along the axis (acrotony). **(c)** Buds before anthesis. **(d)** Frontal view of the flower; the vertical distances from the upper and lower edges of the limb to the throat are roughly comparable. **(e)** Longitudinal dissection of the flower, showing the adaxial structure of the calyx tube and a lateral view of the gynostemium connected to the inferior ovary. **(f)** Top view of the gynostemium, showing the three-lobed stigma during the pistillate phase (female stage) and a unique Ω-shaped margin at each lobe’s junction between the stigma and anthers. **(g)** Lateral view of the flower; the limb reflexes backward and fully covers the calyx tube. **(h)** Leaf surfaces, adaxial (upper) and abaxial (lower): the abaxial side exhibits a faint grayish waxy sheen and a distinct glaucous texture. **(l)** Egg of a birdwing butterfly (*Troides sp.*) laid on the adaxial leaf surface. **(j–m)** Developmental stages from young fruit to seed dehiscence. **(j–l)** Fruit maturation sequence showing progressive enlargement and increasing prominence of longitudinal ridges. **(m)** Seed dehiscence: the pericarp splits via septicidal dehiscence, revealing six distinct carpels that remain apically connected to the central axis. Seeds are tightly attached along each carpel and shed collectively as a unit.

**Type:—** CHINA. Guangxi: Chongzuo City, Longzhou County, Nonggang Village, under tropical evergreen broadleaf forest on karst cliffs, 15 February 2019, Yiwen Jiang (transplanted for cultivation); specimen further collected from the same individual, 22 March 2023, Yifan

Wang & Zirui Guo *yw00080*, leaves, stem & flowers (holotype: IBSC!; isotypes: BAZI!, HITBC!, HK!, IBK!, KUN!, NY!, PE!).

**Paratype:—** CHINA. Guangdong: Guangzhou City, Conghua County, Wenquan Township, Nanxing Village, tropical evergreen broadleaf forest near riparian zone, 12 June 2023, Xiaohui Lin *yw00089* (IBSC!). Guangxi: Baise City, Tiandong County, Naba Township, Fuxing Village, Qipantan Scenic Spot, tropical evergreen broadleaf forest on karst cliffs, 6 May 2024, Yifan Wang & Zirui Guo *VM240501*, leaves, stem, and flowers (IBK!, IBSC!); Nanning City, Wuming District, Damingshan National Nature Reserve, tropical evergreen broadleaf forest on karst cliffs, 3 April 2024, Yundong Huang *yw00060* (IBSC!).

**Diagnosis:—** *Aristolochia magnopurpurea* Y.Fan Wang, Z.R.Guo & J.G.Onyenedum, sp. nov., is morphologically similar to *A. versicolor*, *A. westlandii*, *A. yueguiensis* sp. nov., and *A. pustulata* sp. nov., all characterized by a reflexed calyx limb at anthesis and purple to purplish adaxial limb coloration. However, *A. magnopurpurea* differs from *A. versicolor* in having a much narrower floral throat (ca. 4.2–7.5 mm vs. 12–15 mm) and a uniformly dark purple to blackish-purple, velvety throat and limb surface (vs. yellow, sometimes with minute orange flecks). It is distinguished from *A. pustulata* by the absence of pustulate structures beneath the throat and a smooth, velvety adaxial limb. From *A. yueguiensis*, it differs in both limb and leaf characters: the throat-to-limb margin distances are approximately equal above and below the throat in *A. magnopurpurea* (vs. clearly shorter above the throat in *A. yueguiensis*), and the leaves are elliptic to oblanceolate with a waxy adaxial surface and a distinctly whitish-glaucous abaxial surface (vs. linear leaves with sparse indumentum and no glaucous sheen). Compared to *A. westlandii*, A. magnopurpurea lacks the involute calyx limb and conspicuous white arachnoid venation, instead exhibiting a uniformly reflexed limb with deep purple coloration and no secondary curling.

**Description:—** Liana with twining stems reaching 2–4 m. Roots fusiform or globose, forming a moniliform underground system composed of multiple thickened, tubular segments. Stems terete to oblong-terete; young stems herbaceous, erect, and thinly pubescent; older stems lignified, terete to broadly elliptic in cross-section, with finely fissured, inconspicuous bark. Leaves dimorphic: basal or shade leaves elliptic to ovate; upper or sun-exposed leaves often oblanceolate; apex acuminate; blade 4.0–6.0 × 14.0–20.0 cm; venation pinnate with 9–13 secondary vein pairs; texture membranous to chartaceous. Adaxial surface sparsely pubescent and waxy; abaxial surface with a conspicuous white-glaucous waxy sheen. Petiole 1.5–2.0 cm, thinly pubescent. Inflorescences axillary cymes bearing 1–3 flowers. Peduncle 0.9–1.4 cm, densely rust-villous; pedicels 1.8–4.0 cm, densely brown-villous, twisted. Bracteole solitary, 2.0– 4.0 × 6.0–10.0 mm, densely rust-villous, sessile. Perianth zygomorphic; abaxial surface purple, veined and ridged, rust-villous. Utricle and basal tube not clearly demarcated from the abaxial view; combined length 2.5–3.8 cm. A pale yellow to whitish “window” zone surrounds the calyx-ovary junction on the adaxial side. Adaxial surface of the utricle two-tiered: upper tier dark purple, densely arachnoid-villous, trichomes aligned downward and appressed; lower tier bright red, densely arachnoid-villous, with trichomes loosely ascending and inward-pointing toward the lumen center, often appearing moist and lipidic. Basal tube adaxially purple to dark purple, glabrous or sparsely appressed-pubescent, velvety; narrowing toward the geniculation. Upper tube 1.8–2.8 cm, saccate and swollen near the geniculation; base pale yellow to whitish, sometimes with sparse reddish-purple broken dots; coloration abruptly shifts near the throat to a dark silky purple or purplish-black, velvety and glabrous. Calyx limb 4.5–7.0 cm long, 5.0–7.0 cm in diameter; apex slightly 3-dentate, spreading, strongly reflexed opposite the throat, glabrous, dark purple, velvety. Distance from throat to basal margin approximately equal or slightly shorter to that from throat to proximal margin. Throat subglobose, 4.2–7.5 mm in diameter, slightly ridged at the rim, fleshy, granular-textured; color uniformly velvety purple to purplish-red with occasional white speckling in some populations. Anthers oblong, 2.2–3.5 mm, extrorse. Gynostemium fleshy, 3.5–4.5 mm long, 2.5–5.5 mm in diameter, 3-lobed; each lobe adnate laterally to a granular, omega (Ω)-shaped margin. Ovary inferior, cylindrical, 1.0–1.6 cm long, 6-loculed, with six prominent adaxial ridges, densely rust-villous.

**Distribution and habitat:—** This species is identified in our study as having a strong affinity for karst limestone cliff habitats. Previously misidentified as *Aristolochia versicolor*, it is now recognized as a distinct species occurring primarily in the southern and southwestern regions of Guangxi Zhuang Autonomous Region, with several records extending into adjacent northern Vietnam—areas characterized by typical karst topography (Figure 4e) and a subtropical monsoon climate. We also documented a disjunct population (collectionNo. *yw00089*; Supplementary Table S2) in Guangzhou, Guangdong. Despite this geographic separation, both phylogenetic and morphological evidence confirm that the Guangzhou population belongs to the same species. This disjunction may represent a relict population resulting from ancient geological events or local extinctions, or alternatively, a product of long-distance dispersal. Further fieldwork and population-level studies are required to clarify its biogeographic origin.

**Phenology:—** Flowering from early March to late May; fruiting from late March to mid-June. Seed dehiscence occurs at the basal suture of the capsule approximately two months after pollination. The carpels and seeds detach as a single unit, abscising immediately upon capsule dehiscence.

**Etymology:—** The specific epithet *magnopurpurea* refers to the species’ distinctively large, purple, reflexed calyx limb. Among the reflexed-calyx *Aristolochia* species known to date, including those newly described herein, it bears the second-largest calyx limb, surpassed only by *A. westlandii*, which differs in having conspicuous white venation. The epithet aptly captures this species’ floral morphology.

**Vernacular:—** During our fieldwork, this species—along with several other vegetatively similar *Aristolochia* species—was locally referred to as “⽵叶薯” (zhú yè shŭ) or “⼀拖九⽜” (yī tuō jiŭ niú). To provide a clearer vernacular name that reflects its distinct morphological traits, we propose the Chinese name “⼤紫关⽊通” (dà zǐ guān mù tōng), which conveys both the literal meaning of the epithet and aligns with the commonly used Chinese name for members of subg. *Siphisia*.

**Conservation status:—** *Aristolochia magnopurpurea* has been confirmed as a cross-border species occurring in China and northern Vietnam, with most known populations documented in China. However, the status of populations across the border remains poorly understood. Based on our field investigations, populations are consistently sparse and scattered, typically consisting of fewer than 10 mature individuals. The species is subject to unregulated harvesting of its roots for traditional medicinal use. Alarmingly, during follow-up surveys, we found that approximately half of the populations initially recorded had been excavated and were no longer extant. Germination experiments using collected seeds have been unsuccessful, suggesting potential recalcitrance in seed physiology.

According to the IUCN Red List criteria (2012) and updated guidelines (2024), we identified seven extant populations, each comprising 5–10 mature individuals, with ongoing decline in both population size and habitat quality. The assessed Area of Occupancy (AOO) is 24 km², and the Extent of Occurrence (EOO) is 29,060.81 km². These conditions meet the thresholds for Endangered (EN) under criterion C2a(i), which applies to species with fewer than 2,500 mature individuals, an observed or inferred continuing decline, and no subpopulation containing more than 250 mature individuals. In addition to population decline, we observed that *A. magnopurpurea* serves as a larval host for at least two threatened butterfly species, *Troides aeacus* C. & R. Felder, 1860 and *Troides helena* Linnaeus, 1758 (Figure 7i), both of which were observed ovipositing on this species during fieldwork. The entire genus *Troides*is listed under the Convention on International Trade in Endangered Species of Wild Fauna and Flora (CITES) Appendix II (2024), underscoring its conservation significance. Given the extremely restricted and declining populations, ecological importance, and ongoing anthropogenic pressures, we recommend immediate conservation measures, including local legislative protection, in situ habitat preservation, and comprehensive cross-border field surveys to assess the species’ full conservation status.

*Aristolochia yueguiensis* Y.Fan Wang, Z.R.Guo & J.G.Onyenedum, **sp. nov. Figure 8**.

**Figure 8.**
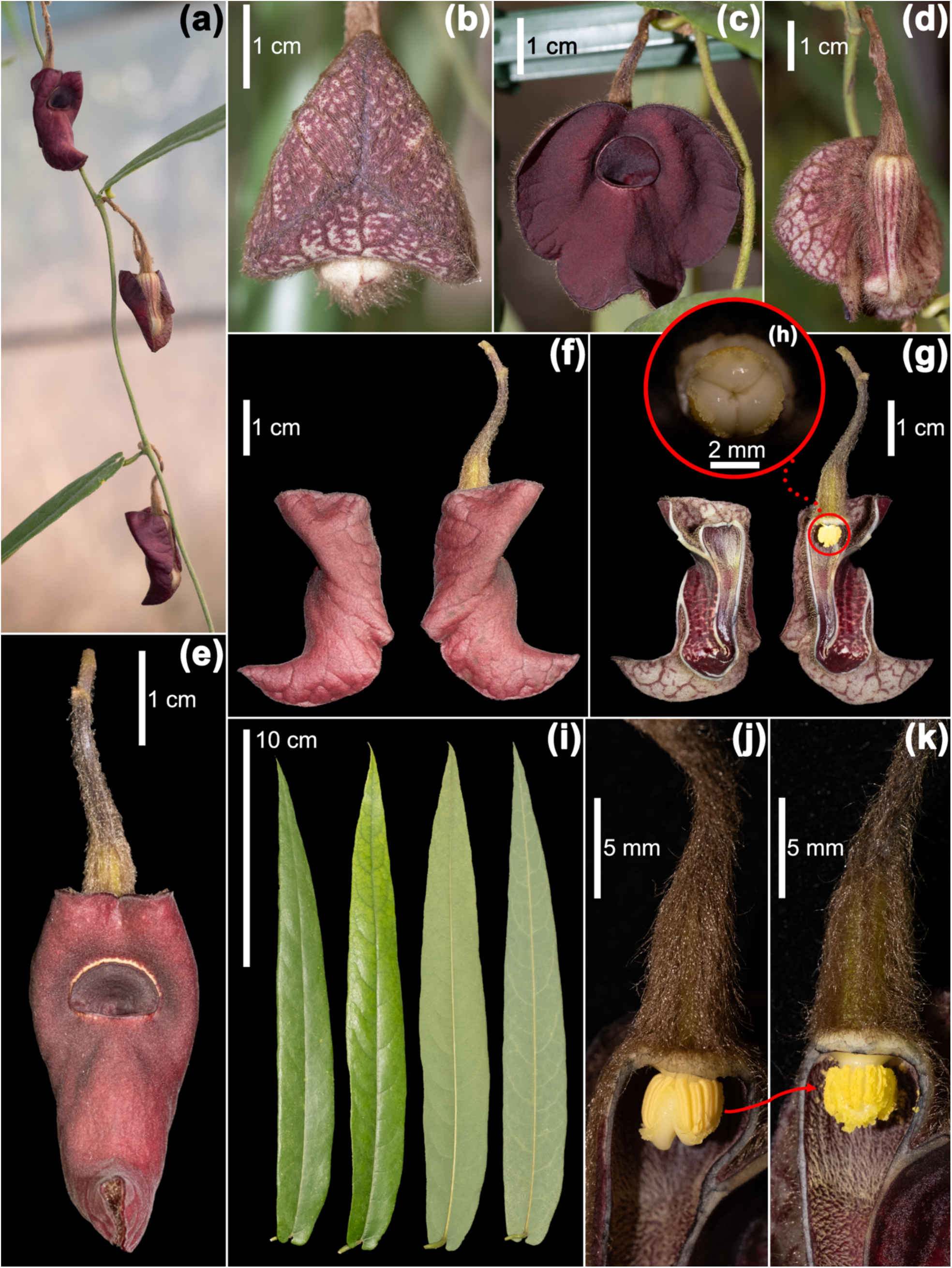
Illustration of *Aristolochia yueguiensis* Y.Fan Wang, Z.R.Guo & J.G.Onyenedum, sp. nov. **(a)** Plant in situ, showing flowering inflorescences axillary from young herbaceous stems. **(b)** Buds before anthesis. **(c)** Frontal view of a freshly opened flower, with the calyx limb not yet reflexed. **(d)** Posterior view of the flower. **(e)** Frontal view of a mature flower with the calyx limb fully reflexed; the vertical distance from the throat to the upper limb edge is much shorter than that to the lower limb edge. **(f)** Lateral view of the flower. **(g)** Longitudinal dissection showing the adaxial structure of the perianth. **(h)** Top view of the three-lobed stigma during the pistillate phase (female stage). **(i)** Leaf blades: two on the left, adaxial surface; two on the right, abaxial surface. (**j–k)** Maturation stages of the gynostemium. **(j)** Pistillate phase: the stigmatic surface is fully exposed and receptive, while the anthers remain tightly appressed to the sides without dehiscence. **(k)** Staminate phase: the stigmatic surface contracts and becomes internalized, while the anthers dehisce and shift downward in position.

**Type:—** CHINA. Guangdong: Zhaoqing City, Fengkai County, Xinghua Township, Shenchong, along riparian zones of stream banks beneath subtropical evergreen broadleaf forests, alt. 317 m, 6 May 2023, Sifeng Chen (transplanted for cultivation); voucher specimens further collected from the same individual, 7 December 2023, Yifan Wang & Zirui Guo *yw00067*, leaves, stem & flowers (holotype: IBSC!; isotypes: BAZI!, HITBC!, IBK!, KUN!, PE!).

**Paratype:—** CHINA. Guangxi: Hechi City, exact locality unknown. Collected from a local seasonal herbal medicine market, sold exclusively as tubular roots. Purchased in June 2023 and subsequently cultivated; voucher specimens further collected from the same individual, 28 December 2023, Yifan Wang & Zirui Guo *yw00079* (IBK!, IBSC!, NY!, US!). Guangxi: Laibin City, Xingbin District, Sanwu Township, Shangyao, growing on slopes under evergreen broadleaf forest, alt. 215 m, 20 May 2024, Haiyu Zhou *VY240501* (BAZI!, CSH!, HK!, HITBC!, IBK!, IBSC!, KUN!, PE!)

**Diagnosis:—** *Aristolochia yueguiensis* Y.Fan Wang, Z.R.Guo & J.G.Onyenedum, sp. nov., resembles *A. westlandii*, *A. magnopurpurea* sp. nov., and *A. pustulata* sp. nov. in having a reflexed, purple adaxial calyx limb, but differs in several key traits. It is readily distinguished from *A. westlandii* by its much smaller flower (calyx 3.2–5.0 × 3.4–4.0 cm vs. 10.0–22.0 × 7.0– 14.5 cm), uniformly dark purple limb lacking white venation, and a limb that reflexes backward and closely appresses the tube without coiling forward—unlike *A. westlandii*, whose limb reflexes then coils forward into an alate form. It differs from *A. magnopurpurea* in the asymmetry of the limb–throat junction: in *A. yueguiensis*, the upper margin lies much closer to the throat than the lower, whereas in *A. magnopurpurea* both distances are roughly equal. Vegetatively, *A. yueguiensis* has linear, leathery, shortly pubescent leaves lacking abaxial glaucous sheen seen in the glabrous, broadly ovate leaves of *A. magnopurpurea*. It is further distinguished from *A. pustulata* by the absence of diagnostic pustulate swellings beneath the throat.

**Description:—** Liana, with twining stems reaching 2–4 m. Roots fusiform to globose, often producing multiple tubular roots arranged in a moniliform underground system. Young stems (1–2 years old) herbaceous, densely rusty-pubescent; older stems woody, compressed, oblong-to elliptic-terete, with fissured bark. Leaf blades narrowly linear, 1.8–5.5 × 15.0–32.0 cm; apex narrowly acute; base small but consistently distinct, cordate or auriculate; venation pinnate, with 11–15 pairs of fine secondary veins; texture leathery. Adaxial surface sparsely rusty-pubescent; abaxial surface pubescent primarily along veins, lamina sparsely pubescent, surface rough-textured. Petioles 1.0–1.5 cm, thinly pubescent, strongly coiled. Inflorescences axillary, cymose, solitary to 3-flowered. Peduncles 0.8–1.0 cm; pedicels 2.4–3.0 cm, densely rusty-villous, twisted. Bracteole solitary per flower, 1.5–2.0 × 2.0–2.5 mm, densely rusty-villous, sessile. Perianth zygomorphic; abaxial surface purple-brown with slightly ridged veins, densely rusty-villous. Utricle and basal tube not clearly distinct abaxially, together 2.5–3.5 cm long to the geniculation; a slightly translucent white window present at the adaxial base surrounding the calyx–ovary junction. Utricle adaxially purple to reddish-purple, densely arachnoid-villous with a whitish covering. Basal tube densely white-villous, purple near the utricle, with villous density gradually decreasing toward the geniculation; tube narrowing markedly toward the geniculation. The upper tube 2.2–3.0 cm long, inflated near the geniculation, forming a secondary saccate swelling; adaxially whitish with purple spots, transitioning toward the throat into a purple-reddish tone with increasingly dense spotting—sometimes coalescing into a uniform purple surface. Surface velvety, glabrous. Calyx limb 3.2–5.0 cm long, 3.4–4.0 cm in diameter, 3-dentate, reflexed opposite the throat, glabrous, dark purple, velvety; distance from throat to basal limb margin greater than that to the apical margin; inferior lobe recurved, forming an upturned, hook-like structure. Throat arcuate to subglobose, 6–13 mm in diameter, varying across individuals and populations, dark purple, velvety, slightly ridged along the rim, fleshy, smooth. Anthers oblong, 1.8–2.7 mm, extrorse. Gynostemium fleshy, 3.5–5.0 mm long, 3.0–4.8 mm in diameter, 3-lobed; apex obtuse, each lobe laterally adnate to a granular margin. Ovary inferior, cylindric, 0.9–1.5 cm, 6-locular, with six prominent ridges, surface densely rusty-villous.

**Distribution and habitat:—** This species is typically found at low elevations on loess hills in western Guangdong Province and the northern part of Guangxi Zhuang Autonomous Region. It inhabits subtropical monsoon evergreen broadleaf forests, growing near water sources such as streams and waterfalls, often in rocky microhabitats (Figure 4e).

**Phenology:—** Flowering from late February to June, with variation depending on locality; fruiting from March to mid-July. Seeds are retained within the carpophore and detach as the capsule dehisces septicidally from the basal end (proximal to the pedicel). The carpophore abscises within a day of dehiscence, falling as a unit with seeds still attached to the carpels.

**Etymology:—** The epithet *yueguiensis* was selected to emphasize the interprovincial distribution of this species between Guangdong Province (abbreviated in Chinese Pinyin as “Yue”) and Guangxi Province (“Gui”). This species occurs naturally in both provinces, and the epithet reflects its occurrence in these distinct but geographically contiguous regions.

**Vernacular:—** Locally, this plant is referred to as “⽵叶薯” (zhú yè shŭ) or “⼀拖九⽜” (yī tuō jiŭ niú), names also applied to *A. magnopurpurea*, *A. championii*, and other sympatric species with similar moniliform roots, which are often harvested as herb medicine and traded regionally under these umbrella terms. To distinguish this species and reflect its etymological and geographical identity, we propose the Chinese vernacular name “粤桂关⽊通” (yuè guì guān mù tōng).

**Conservation status:—** *Aristolochia yueguiensis* was documented by our study across four populations in southern China, specifically in western Guangdong Province and northern to eastern Guangxi Zhuang Autonomous Region, with highly variable population densities. In most cases, the species was observed as a solitary population comprising only one to five mature individuals. Despite repeated field visits revealing flowers, mature capsules, and abundant seeds, juvenile individuals were rarely encountered. Concurrently, this species is under increasing pressure from unregulated harvesting, as its roots are sought after for use in traditional medicine. Several populations initially documented could not be relocated in subsequent surveys, likely due to extraction for medicinal purposes. This species was also frequently encountered in informal, unregulated herbal markets across Guangdong and Guangxi, where roots and lignified stems (identity confirmed through cultivation) were sold under general vernacular names applied to multiple *Aristolochia* species.

Based on our assessment, wild populations likely comprise fewer than 2500 mature individuals in total. Considering the observed rapid decline, ongoing overexploitation, and highly fragmented and sparse distribution, and following the IUCN Red List criteria (2012) and updated guidelines (2024), we propose classifying *Aristolochia yueguiensis* as Endangered (EN) under criterion C2a(i), which applies to species with fewer than 2,500 mature individuals, a continuing decline in numbers, and no subpopulation exceeding 250 individuals. Its urgent conservation need is underscored by both natural vulnerability and anthropogenic threats.

*Aristolochia pustulata* Y.Fan Wang, Z.R.Guo & J.G.Onyenedum, **sp. nov. Figure 9**.

**Figure 9.**
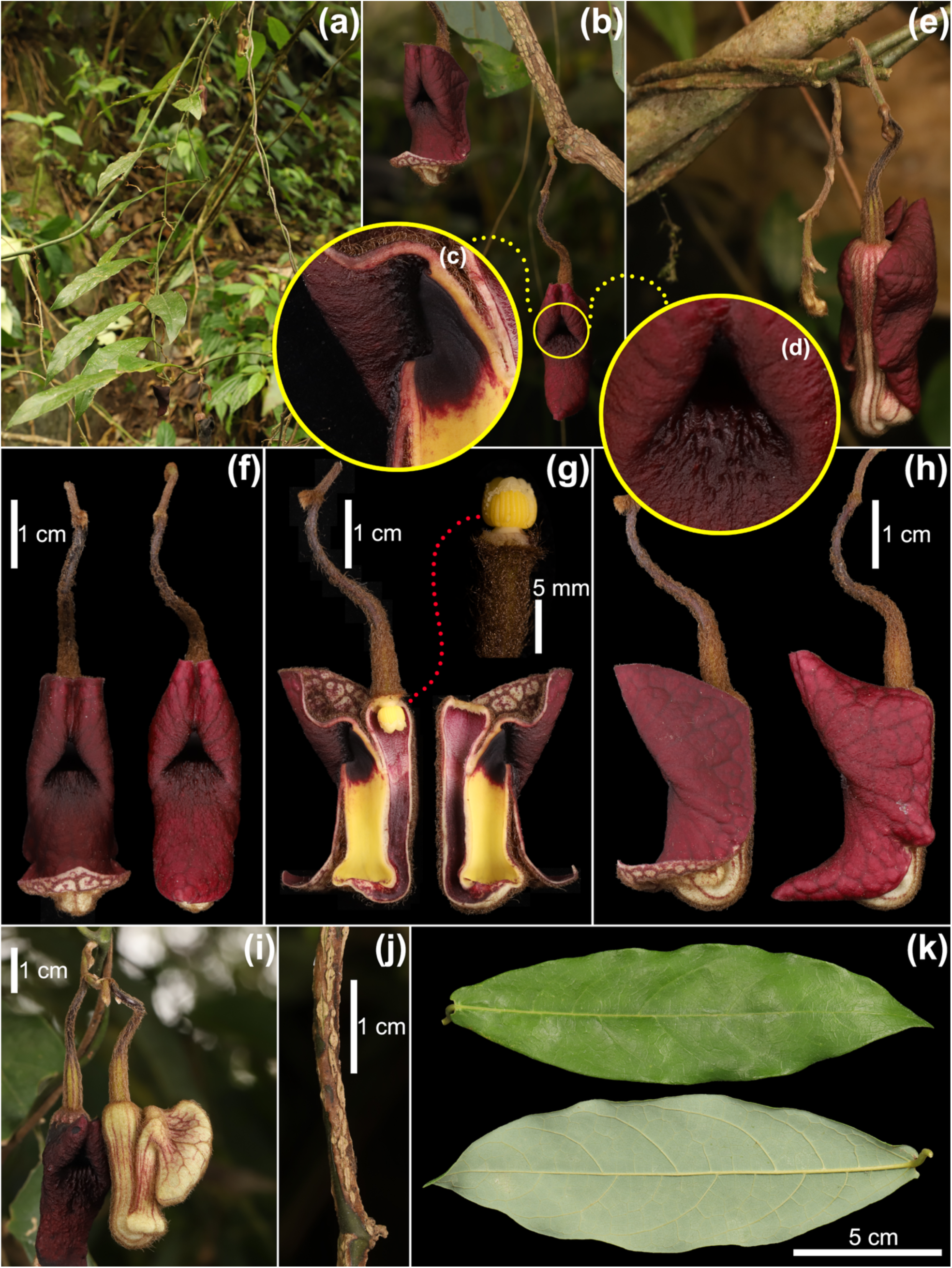
Illustration of *Aristolochia pustulata* Y.Fan Wang, Z.R.Guo & J.G.Onyenedum, sp. nov. **(a)** Plant in situ, showing flowering inflorescences axillary from young herbaceous stems. **(b)** Cauliflorous inflorescence arising from dormant or latent meristems on lignified old stems. **(c– d)** Close-up of pustulate structures at the calyx throat entrance: **(c)** Concentration of dark purple pigmentation near the adaxial connection between the upper tube and the throat; **(d)** Papillate pustules distributed on the calyx limb just beneath the throat, representing the most diagnostic feature of this species. **(e)** Posterior view of the flower. **(f)** Frontal view of the flower. **(g)** Longitudinal dissection showing the adaxial surface of the calyx tube and a lateral view of the gynostemium. **(h)** Lateral view of the flower. **(i)** Buds in situ; the RICH syndrome is already conspicuous before anthesis, as indicated by the prominent inflation of the basal tube. **(j)** Lignifying old stem, showing the development of coarse corky bark from the originally herbaceous green stem. **(k)** Leaf, showing both adaxial (upper) and abaxial (lower) surfaces.

**Type:—** CHINA. Yunnan: Wenshan Zhuang and Miao autonomous prefecture, Malipo county, Xiajinchang Township, Yanfeng cave, on the slopes of karst limestone mountains under subtropical forest, alt. 1321 m, 17 March 2024, Yifan Wang, Zirui Guo & Qing Yu *yw00081*, leaves, stem & flowers (holotype: IBSC!; isotypes: BAZI!, CSH!, HK!, HITBC!, IBK!, KUN!, NY!, PE!, US!).

**Diagnosis:—** *Aristolochia pustulata* Y.Fan Wang, Z.R.Guo & J.G.Onyenedum, sp. nov., is morphologically similar to *A. magnopurpurea* sp. nov. and *A. yueguiensis* sp. nov., all characterized by a reflexed calyx limb uniformly purple on the adaxial surface. However, *A. pustulata* is readily distinguished by the unique presence of conspicuous pustules on the calyx limb beneath the throat—a feature not reported in any other *Aristolochia* species. Furthermore, in *A. pustulata*, the calyx limb forms prominent folds that rise up and partially obscure the throat from a frontal view, creating a distinctive triangular pseudo-throat. This combination of pustulate ornamentation and obscured throat structure makes *A. pustulata* easily diagnosable from the aforementioned congeners.

**Description:—** Liana, reaching up to 3–5 m. Roots fusiform or globose, often producing multiple tubular roots arranged in a moniliform underground system. Stems terete to oblong-terete; young stems herbaceous, thinly pubescent; older stems lignified, terete or broadly elliptic in cross-section, with finely fissured bark. Leaf blades obovate-elliptic to oblanceolate, 3.0–6.0 × 13.0–21.5 cm; apex acuminate; venation pinnate, with 9–13 pairs of secondary veins; texture membranous to coriaceous. Adaxial surface sparsely pubescent to glabrous; abaxial surface glabrous. Petioles 1.5–1.8 cm, thinly pubescent, twisted. Inflorescence axillary, cymose, each cyme bearing 1–5 flowers; flowers open basipetally. Peduncles 0.8–1.4 cm, rusty-villous. Pedicels 1.8–2.0 cm, densely brown-villous, twisted. Bracteole one per flower, 1.1–1.3 × 1.8–2.0 mm, densely rusty-villous. Perianth zygomorphic; abaxial surface brown-whitish with ridged purplish venation, golden to rusty-villous. Utricle and basal tube not clearly distinct externally, together 3.5–3.8 cm down to the geniculation; a translucent white window is present adaxially at the calyx–ovary junction. The adaxial utricle surface is two-tiered: the upper tier purple to red, densely arachnoid-villous with white appressed trichomes; the lower tier bright pinkish-red, lipidic and waxy, with upright trichomes pointing toward the tube, densely white-villous. Basal tube purple to dark purple, with a short but dense white indumentum and velvety texture; tube narrows and becomes constricted toward the geniculation. Upper tube 1.8–2.5 cm, inflated at the geniculation, forming a secondary spherical saccate structure with purple dots adaxially surrounding the basal tube opening. The inflated surface is bright yellow at lower portion, abruptly transitioning to dark purple or black on the upper portion near the throat; surface silky, velvety, and glabrous. Calyx limb adaxially 2.8–4.5 cm long, 3.0–4.8 cm in diameter, 3-dentate at apex, spreading and reflexed opposite the throat; glabrous, reddish-purple, leathery. The upper lateral lobes rise prominently when reflexed, forming a triangular pseudo-throat structure that partially obscures the true calyx throat. Calyx throat semicircular, 4.0–5.8 mm in diameter, dark purplish, not fully visible from the exterior. Beneath the throat on the limb, a dark purple muricated region bears fleshy, upward-pointing pustules. Anthers oblong, 2.9–3.1 mm, extrorse. Gynostemium fleshy, 4.0–5.0 mm long, 2.8–3.5 mm in diameter, 3-lobed; apex obtuse, each lobe laterally adnate to a granular margin. Ovary inferior, cylindric, 1.0–1.5 cm, with six adaxial ridges obscured beneath dense rusty villous indumentum.

**Distribution and habitat:—** *Aristolochia pustulata* is a highly endemic species, currently known only from Wenshan Zhuang and Miao Autonomous Prefecture, specifically in Xichou and Malipo counties (Figure 4e), where it inhabits karst mountains under subtropical monsoon climate and cloud forest conditions. While such narrow and fragmented distributions are common among *Aristolochia* species, the pattern observed here likely reflects strong ecological specialization and localized speciation driven by microhabitat conditions. The known populations are located near the China–Vietnam border, where geopolitical boundaries do not reflect discontinuities in geological or bioclimatic conditions. Although this study did not include surveys on the Vietnamese side, we hypothesize that *A. pustulata* may also occur in adjacent regions of northern Vietnam with similar environmental settings.

**Phenology:—** Flowering occurs from early March to late April; fruiting from late March to mid-May. Seeds are retained within the carpophore and are released as the capsule dehisces septicidally from the basal end (proximal to the pedicel). The carpophore abscises within a day of dehiscence, falling as a unit with seeds still attached to the carpels.

**Etymology:—** The epithet *pustulata* refers to the species’ distinctive morphological feature near the calyx throat, a darkened purple, muricate region bearing fleshy, upward-pointing pustules, not observed in any other known *Aristolochia* species.

**Vernacular name:—** This species has no known local name, as confirmed by our field surveys. To support future efforts in conservation and ethnomedicinal research, we propose the Chinese vernacular name “ 棘喉关⽊通” (jí hóu guān mù tōng), reflecting the diagnostic pustulate structures beneath the calyx throat.

**Conservation status:—** *Aristolochia pustulata* is currently known only from Wenshan Zhuang and Miao Autonomous Prefecture in southeastern Yunnan, with over eight populations documented: three in Xichou County and five in Malipo County. Each population comprises approximately 5 to 20 mature individuals, all observed to be flowering and reproductively mature. Fruit capsules were also recorded, and seedling presence suggests relatively stable population structure at some sites. However, the species appears to exhibit a high degree of endemism and is narrowly restricted to these localized karst habitats, indicating ecological specialization to specific microhabitats.

Although no direct evidence of medicinal exploitation has been identified, *A. pustulata* is threatened by habitat loss due to expanding agricultural activities. In particular, we observed corn (*Zea mays* L.) and tsaoko cardamom (*Amomum tsao-ko*) plantations directly adjacent to several populations. Land clearance for such cultivation typically results in complete habitat removal. The calculated Area of Occupancy (AOO) is 48.000 km², and the Extent of Occurrence (EOO) is 1,448.408 km². According to the IUCN Red List Categories and Criteria (2012) and the latest Guidelines for using it (2024), these values meet the thresholds under criterion B1ab(iii)+2ab(iii). Therefore, we propose *A. pustulata* be classified as Endangered (EN) based on its restricted distribution, ecological specialization, and ongoing habitat degradation.

*Aristolochia nitida* Y.Fan Wang, Z.R.Guo & J.G.Onyenedum, **sp. nov. Figure 10**.

**Figure 10.**
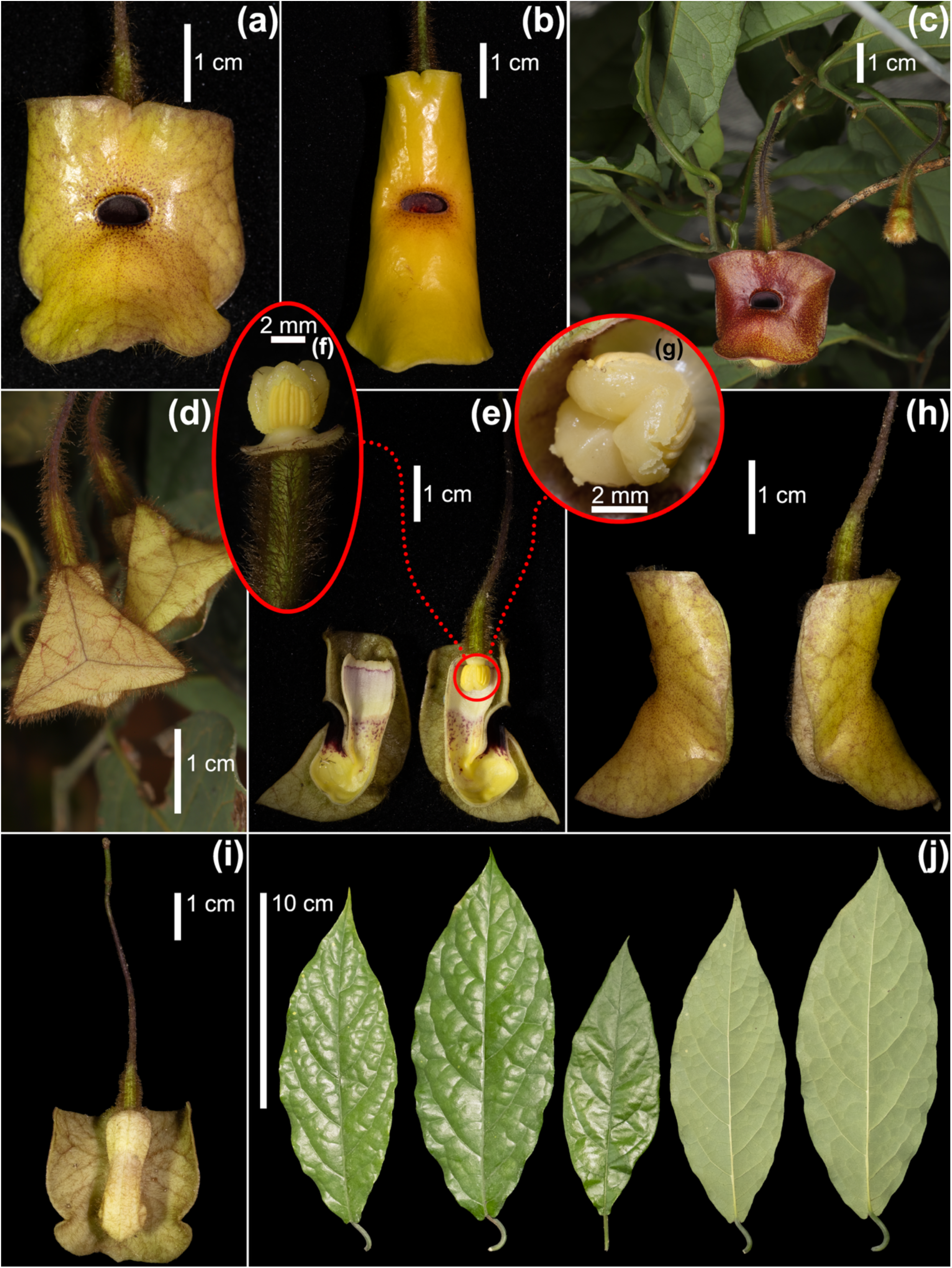
Illustration of *Aristolochia nitida* Y.Fan Wang, Z.R.Guo & J.G.Onyenedum, sp. nov. **(a–c)** Frontal view of flowers, showing intraspecific variation in calyx limb pigmentation among three individuals. Variation in adaxial pigmentation is widely observed both among individuals and among different flowers on the same individual, indicating that calyx limb color is not a stable or reliable diagnostic trait for identification. Slight shape variation in the calyx throat is also observed; while generally elliptical, minor differences occur between individuals and between flowers on the same plant. **(c)** additionally shows a flower in situ. **(d–j)** Floral and vegetative morphology based on an individual corresponding to floral type **(a)**. **(d)** Bud, abaxially rusty-villous. **(e)** Longitudinal dissection of the flower, showing adaxial floral structure; the “RICH” syndrome is present but less pronounced in *A. nitida*, particularly the basal tube constriction, which is more developed in other members of the *A. versicolor* species group. **(f)** Lateral view of the gynostemium. **(g)** Top view of the gynostemium. **(h)** Lateral view of the flower. **(i)** Posterior view of the flower. **(j)** Leaf blades, adaxial (left three) and abaxial (right two) surfaces, showing conspicuous bullate (blistered) venation.

**Type:—** CHINA. Yunnan: Wenshan Zhuang and Miao autonomous prefecture, Malipo county, Babu township, Nadeng village, under subtropical cloud forest on slopes of limestone karst mountains, alt. 1470 m, 8 July 2023, Fuguan Deng (transplanted for cultivation); specimens further collected from the same individual, 21 March 2024, Yifan Wang & Zirui Guo *yw00082*, leaves, stems & flowers (holotype: IBSC!; isotypes: BAZI!, CSH!, HK!, HITBC!, IBK!, KUN!, NY!, PE!, US!).

**Paratype:—** CHINA. Yunnan: Honghe Hani and Yi Autonomous Prefecture, Hekou Yao Autonomous County, Nanxi Township, 2018, Lei Cai (transplanted for cultivation at Kunming Institute of Botany); voucher specimen later collected from the same individual, 25 March 2025, Zirui Guo *yw00091*, leaves, stems & flowers (HITBC!, KUN!); Wenshan Zhuang and Miao Autonomous Prefecture, Malipo County, Babu Township, Nadeng Village, under subtropical cloud forest on slopes of limestone karst mountains, alt. 1271 m, 20 April 2024, Fuguan Deng (transplanted for cultivation); specimens further collected from the same individual, 29 October 2024, Zirui Guo *202410290001* (IBK!, KUN!).

**Diagnosis:—** *Aristolochia nitida* Y.Fan Wang, Z.R.Guo & J.G.Onyenedum, sp. nov., resembles *A. versicolor* and *A. longii* sp. nov. in possessing a reflexed yellow to reddish-orange adaxial calyx limb. However, it is readily distinguished by the smoothly continuous transition between the basal and upper perianth tubes, marked only by a slight constriction—unlike the pronounced narrowing of the basal tube seen in most reflexed-limb species. The adaxial tube is uniformly bright yellow and unspotted, while the adaxial limb surface is glossy and lustrous, in contrast to the velvety texture of *A. longii*. It also differs in limb size, being significantly larger (2.7–4.6 × 2.7–5.6 cm) than that of *A. longii* (1.6–2.7 × 1.6–2.7 cm). Compared to *A. versicolor*, which typically has greenish-yellow limbs with occasional purplish hues and a wide, open throat (1.0–1.4 cm) often marked by scattered orange flecks, *A. nitida* is characterized by a consistently broad, elliptic, flat calyx throat, dark purplish-red in color and measuring 4.0–8.5 mm in diameter. Very noticeably, this species exhibits variation in calyx limb coloration, ranging from pure bright yellow to yellow with fine brown speckles, and in some cases, orange (Figure 10a–c, showing three floral forms). Therefore, limb color alone should not be relied upon as a diagnostic character for species identification.

**Description:—** Liana, reaching up to 2–2.5 m. Roots fusiform or globose, with swollen tuberous segments connected in series, forming a moniliform geophytic system. Stems terete; young stems herbaceous, thinly golden-pubescent; older stems lignified, terete, reaching up to 4 cm in diameter, with a yellowish central radiating medullary ray. Leaf blades elliptic to ovate, 4.0–6.0 × 11.0–14.0 cm; apex abruptly acuminate or caudate; venation pinnate, with 6–8 pairs of secondary veins; texture papery and rugose. The lamina between veins is distinctly convex on the adaxial side, producing a bubbled, wrinkled texture. Both surfaces glabrous. Petioles 1.6–2.5 cm, thinly pubescent. Inflorescences axillary, cymose, with each cyme bearing 1–4 flowers. Pedicels 2.3–4.5 cm, short-pubescent, dark purplish near the axil, becoming green toward the ovary, twisted. Bracteole solitary per flower, 1.1–1.5 × 2.0–2.2 mm, densely rusty-villous, sessile. Perianth zygomorphic; abaxial surface light yellow, evenly covered in long rusty villous hairs; venation on the abaxial limb inconspicuous. Utricle and basal tube not clearly demarcated externally, together 2.0–3.5 cm to the geniculation; a translucent whitish window pane is present adaxially around the calyx–ovary junction. Adaxial utricle surface two-tiered: upper tier red-purple with dense, short, appressed white indumentum; lower tier pinkish-white with dense, erect white indumentum. Basal tube adaxial surface bright yellow, waxy, glossy, and glabrous, showing only slight constriction and merging smoothly into the upper tube. Geniculation maintains a continuous, smooth transition with consistent yellow, glabrous, waxy texture. Upper tube 1.3–2.3 cm, transitioning abruptly near the throat into a velvety dark purple or black surface adaxially. Calyx limb 2.7–4.6 cm long, 2.5–5.8 cm in diameter, 3-dentate at the apex, spreading or reflexed opposite the throat; The degree of limb reflexion varies among flowers, individuals, and populations. In some individuals, the lobes are fully reflexed and closely appressed to the tube; in others, the lobes are coplanar, forming a conspicuous orbicular, disk-shaped limb. Limb surface glabrous, consistently waxy, glossy, and lustrous; coloration ranges from bright yellow to orange-red, often with varying densities of fine yellow speckles. Color variation occurs both among and within populations. The distance from the throat to the basal margin exceeds that to the proximal (upper) margin. Throat broadly oval to elliptic, 4.0–8.5 mm in diameter, laterally broader than tall, purplish-red, velvety, and glabrous. Anthers oblong, 1.9–4.2 mm, extrorse. Gynostemium fleshy, 4.0–6.0 mm long, 3.0–5.8 mm in diameter, 3-lobed; lateral sides of each lobe adnate to a granular margin. Ovary inferior, cylindric to conical, 1.3–1.7 cm, densely rusty-villous on the abaxial surface, with six ridges.

**Distribution and habitat:—***Aristolochia nitida* is currently known from karst limestone mountains with cloudy, evergreen subtropical forests. Populations are scattered along the China–Vietnam border, ranging from Napo County in Baise, Guangxi, to Hekou County in Honghe, Yunnan (Figure 4e). The species typically grows in rocky substrates, anchoring itself with fusiform and swollen roots in areas where the humus layer is thin. Many of the documented populations are located within 50 km of the international border. Despite this geopolitical boundary, the climatic and ecological conditions across the region are continuous. We therefore hypothesize that additional populations may exist in adjacent areas of northern Vietnam.

**Phenology:—** Flowering from late March to mid-June; fruiting from mid-April to early July.

**Etymology:—** The epithet *nitida* is derived from the species’ distinctive feature—a lustrous, waxy adaxial calyx limb. Its exceptionally glossy surface renders the species unique and readily distinguishable from other *Aristolochia* species.

**Vernacular name:—** This species has no known vernacular name and is not recognized by the local community. Based on its etymology, we propose the Chinese vernacular name “光泽关⽊通” (guāng zé guān mù tōng), which accurately reflects its defining glossy characteristic.

**Conservation status:—** This border-distributed species is currently known from only four populations, each comprising approximately 20 mature individuals. Within each population, we observed both fruiting individuals and substantial numbers of seedlings, suggesting ongoing recruitment. However, outside the karst cloud forests along the China– Vietnam border, no additional populations have been identified, indicating a high degree of habitat specificity and endemism. Although no medicinal use of this species was reported among local communities, its habitat is increasingly threatened by agricultural expansion. We observed corn (*Zea mays*) and tsaoko cardamom (*Amomum tsao-ko*) plantations in close proximity to several populations, indicating active encroachment into suitable habitat.

Based on the populations documented in this study, the Area of Occupancy (AOO) is 20.000 km² and the Extent of Occurrence (EOO) is 2,530.857 km². Following the IUCN Red List Categories and Criteria (2012) and the updated Guidelines (2024), *Aristolochia nitida* qualifies for listing as Endangered (EN) under criterion B1ab(iii)+2ab(iii). This reflects its limited and fragmented distribution (EOO < 5,000 km²; AOO < 500 km²), presence at fewer than 10 locations, and an inferred continuing decline in habitat quality due to ongoing agricultural pressure.

*Aristolochia longii* Y.Fan Wang, Z.R.Guo & J.G.Onyenedum, **sp. nov. Figure 11**.

**Figure 11.**
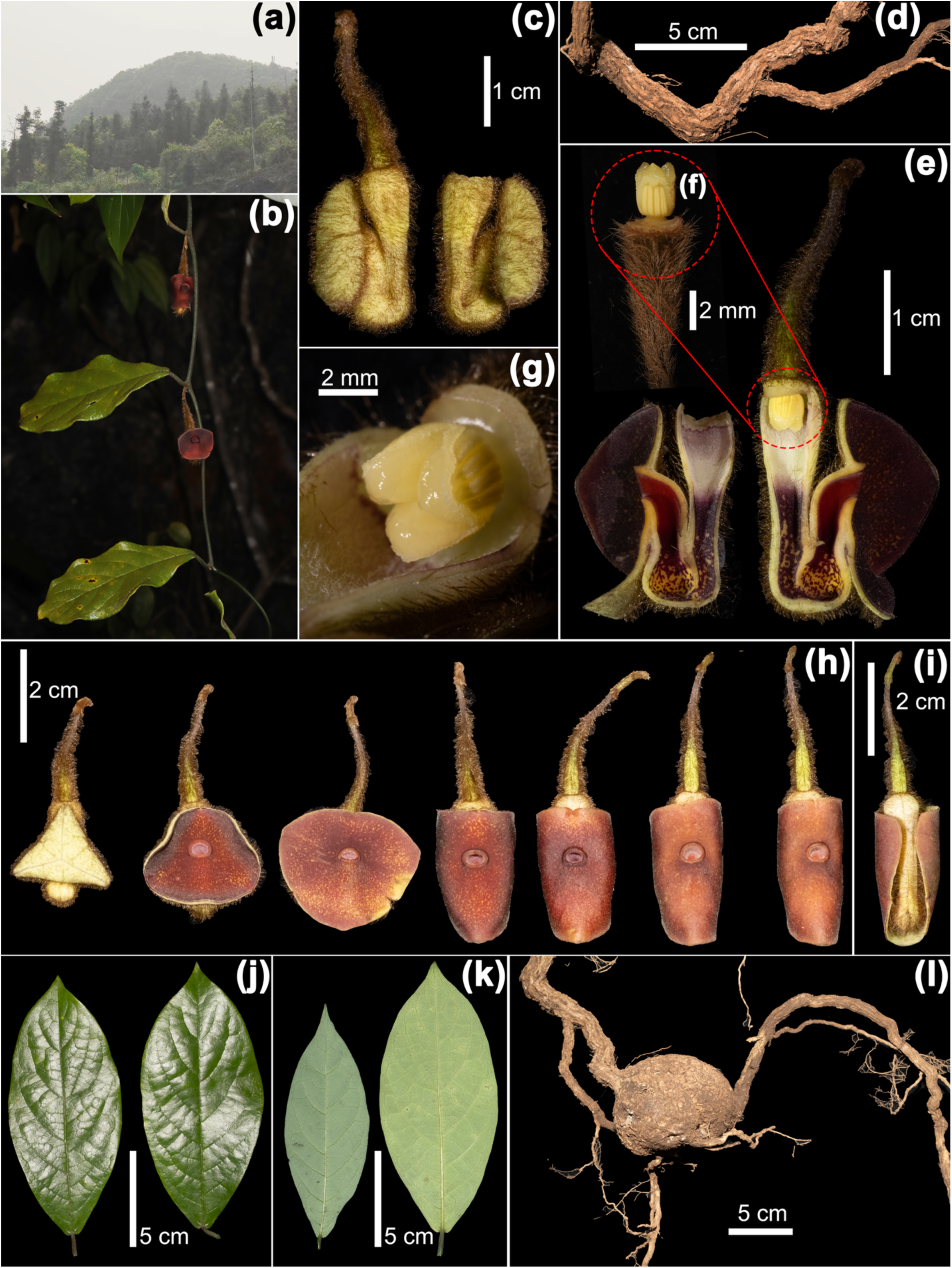
Illustration of *Aristolochia longii* Y.Fan Wang, Z.R.Guo & J.G.Onyenedum, sp. nov.**(a)** Habitat: karst limestone cloud forest in southwestern China. **(b)** Plant flowering in situ, typically bearing solitary axillary inflorescences. **(c)** Abaxial view of the bud. **(d)** Lignified old basal stem. **(e)** Longitudinal dissection of the flower, showing adaxial structures. **(f)** Lateral view of the gynostemium connected to the ovary. **(g)** Close-up of the three-lobed stigma during the pistillate phase (female stage). **(h)** Stages of calyx limb development post-anthesis: the three-dentate calyx limb progressively rolls back to eventually cover the perianth tube. **(i)** Posterior view of the flower. **(j)** Leaf, adaxial surface. **(k)** Leaf, abaxial surface. **(l)** Fusiform and moniliform tubular roots.

**Type:—** CHINA. Yunnan: Wenshan Zhuang and Miao autonomous prefecture, Malipo county, Tianbao Township, Tianbao Village, roadside from Diaozhuping to Muliangzhai, growing in crevices of karst limestone, alt. 1372 m, 16 March 2024, Yifan Wang, Zirui Guo & Qing Yu *yw00087*, leaves, stems & flowers (holotype: IBSC!; isotypes: BAZI!, HITBC!, IBK!, KUN!, NY!, PE!, US!).

**Paratype:—** CHINA. Yunnan: Wenshan Zhuang and Miao autonomous prefecture, Malipo county, Tianbao Township, Tianbao Village, Diaozhuping, growing in crevices of karst limestone, alt. 1381.3 m, 15 April 2024, Fuguan Deng *yw00092*, leaves, stems & flowers (HITBC!, KUN!).

**Diagnosis:—** *Aristolochia longii* Y.Fan Wang, Z.R.Guo & J.G.Onyenedum, sp. nov., is morphologically similar to *A. nitida* sp. nov., with both species bearing elliptic to ovate leaves and a reflexed calyx limb. However, *A. longii* is readily distinguished by its significantly smaller perianth—the smallest among all known calyx-reflexed species—with the calyx measuring only 1.6–2.7 × 1.6–2.7 cm (vs. *A. nitida* at 2.7–4.6 × 2.5–5.6 cm). In *A. longii*, the calyx limb is velvety adaxially, colored purplish-red to brownish-red, and the perianth structure shows a clearly constricted basal tube and an inflated upper tube—features that are only weakly expressed in *A. nitida*. Vegetatively, *A. longii* has thin, smooth, chartaceous leaves with a flat surface, in contrast to the distinctly blistered texture of *A. nitida*’s foliage.

**Description:—** Liana with vines reaching up to 2 m. Roots fusiform or globose, often producing multiple tubular roots arranged in a moniliform underground system. Stems terete to oblong-terete; young stems herbaceous, thinly pubescent; older stems lignified, terete, with fissured bark. Leaf blades elliptic to ovate, 4.0–8.5 × 9.0–18.0 cm; apex acuminate or apiculate; venation pinnate with 8–10 pairs of secondary veins; texture chartaceous. Adaxial surface smooth, glabrous, and slightly lustrous; abaxial surface glabrous except for sparse indumentum along the veins. Petioles 1.0–2.2 cm, thinly covered with short indumentum, twisted. Inflorescences axillary, each cyme typically bearing a solitary flower, occasionally up to three; however, only the terminal flower usually develops fully, while the others remain undeveloped. Pedicels 0.7–1.5 cm, densely rusty-villous, twisted. Bracteole solitary, 1.1–1.5 × 2.0–2.2 mm, densely rusty-villous, sessile. Perianth zygomorphic; abaxial surface bright yellow-green, smooth, velvety, and thinly rusty-pubescent. Utricle and upper tube together 2.2–2.5 cm, with no distinct boundary between them. A translucent whitish window is present adaxially around the calyx– ovary junction. The adaxial utricle surface is two-tiered: upper tier purplish-pink, lower tier white; both tiers covered with short tomentum. The upper tube transitions from purple with short tomentum to a more glabrous purplish surface with yellow patches and sparse hairs, ultimately becoming velvety. It gradually narrows toward the geniculation. Upper tube 1.1–1.6 cm, forming a secondary saccate swelling at its junction with the basal tube; this inflated area appears orange-yellow with purple patches. Toward the throat, the tube transitions from purple to reddish-purple, velvety, and pubescent. Calyx limb 1.6–2.7 cm long, 1.6–2.7 cm in diameter, 3-dentate, spreading and reflexed opposite the throat, appressed to the tube; adaxial surface glabrous, purplish-red, and velvety. The distance from the throat to the basal margin is approximately equal to, or slightly greater than, that to the proximal margin. Calyx throat circular, 3.0–5.4 mm in diameter, purplish-red with small yellow patches or entirely purplish; sparsely pubescent with short indumentum; slightly ridged along the rim; inner surface smooth. Anthers oblong, 1.5–2.7 mm, extrorse. Gynostemium fleshy, 3.0–3.5 mm long and in diameter, 3-lobed; apex obtuse; lateral sides of each lobe adnate to a granular margin. Ovary inferior, cylindric to conical, 1.0–1.5 cm long, 6-locular, with six prominent adaxial ridges; abaxial surface densely rusty-villous.

**Distribution and habitat:—** *Aristolochia longii* occurs in cloud forests on karst limestone at elevations of 1200–1400 m and shares a close geographic distribution with *A. pustulata* and *A. nitida*, both of which are also found in Malipo County. To date, only two populations have been discovered, both located in Tianbao Township, Malipo County, Wenshan, Yunnan (Figure 4e). *A. longii* appears to be a highly endemic species, restricted to this narrowly specialized ecological niche. In its natural habitat, the geophytic root system forms swollen tubers that anchor the plant directly into limestone crevices, often in shallow humus-rich leaf litter. The habitat is characterized by persistent humidity and dense fog throughout much of the year, with ambient moisture frequently exceeding 90% on foggy days. These microclimatic and edaphic conditions likely contribute to the species’ high degree of endemism and limited distribution.

**Phenology:—** Flowering from mid-March to late April; fruiting and seed development not yet observed.

**Etymology:—** The epithet *longii* honors Mr. Mingfeng Long, who first discovered this plant and has made a profound impact on plant survey and conservation efforts in the Wenshan Zhuang and Miao autonomous prefecture, Yunnan through his dedication to botanical work.

**Vernacular name:—** This species does not have a known vernacular name. We propose the Chinese vernacular name “龙⽒关⽊通” (lóng shì guān mù tōng), which accurately reflects the etymological dedication of the species.

**Conservation status:—** To date, we have documented only two populations of this species within an extremely restricted region near the Chinese–Vietnamese border. This area lacks formal conservation status and is subject to severe pressures from agricultural expansion and grazing. One population consists of a single, declining individual directly exposed at the roadside, exhibiting signs of poor health. The second population occurs in a relatively undisturbed montane site, with approximately 20 individuals, including both mature and juvenile plants, indicating a stable population structure. For conservation purposes, the type individual—belonging to the first, vulnerable population—was transplanted and propagated through cuttings distributed to several trusted scientific and educational institutions for ex situ preservation. The plant has since acclimated to cultivation and shown signs of recovery.

Despite extensive surveys in the region and in ecologically similar habitats, no additional populations have been located. Based on the IUCN Red List Categories and Criteria (2012) and the updated Guidelines (2024), *Aristolochia longii* qualifies as Critically Endangered (CR) under criterion D, which applies to species with fewer than 50 mature individuals. This designation reflects its extremely limited distribution, lack of habitat protection, and small population size confined to a cross-border region under active human disturbance.

## 6. Conclusion

This study presents a well-resolved phylogenomic framework for *Aristolochia* subg. *Siphisia*, incorporating broad taxonomic coverage, including numerous previously unsampled taxa. Our results provide species-level resolution and support a seven-clade systematic delineation grounded in molecular evidence, biogeographic boundaries, and shared developmental synapomorphies. These findings bring long-overdue clarity to *Siphisia*, a lineage historically understudied or overshadowed by the canonical *Aristolochia* s.s. group.

We also assessed the extent of introgression and incomplete lineage sorting (ILS) within *Siphisia*. While hybridization signals were minimal, ILS emerged as the primary driver of gene– species tree discordance. This widespread ILS signal, together with short branch lengths and lineage-specific morphological traits, supports a possible scenario of rapid speciation across the clade. In parallel, our observations of floral architecture, particularly the absence of the trap-and-release mechanism found in *Aristolochia* s.s., suggest that *Siphisia* may have evolved alternative pollination strategies. Building on our robust systematic foundation, we revisited the long-standing taxonomic uncertainty surrounding *Aristolochia versicolor*. We resolved its phylogenetic position and morphological confusion, identified and described five new species, and defined a monophyletic species group characterized by a hypothetical pollinator filtering mechanism shaped by the RICH syndrome. This strategy likely reflects ecological adaptation and may function to reinforce reproductive isolation through targeted pollinator selectivity.

Future work should expand taxon sampling to include the rapidly growing number of newly described or unsequenced species and employ more refined phylogenomic methods. Such efforts will help complete the evolutionary portrait of *Siphisia* and contribute to a broader understanding of diversification patterns in Magnoliids and angiosperms at large.

## Declaration of Generative AI and AI-Assisted Technologies in the Writing Process

During the preparation of this work, the authors used the ChatGPT-4o model solely for grammar and language refinement. After employing this tool, the authors reviewed and edited the content as needed and take full responsibility for the final version of the manuscript.

## Supplementary Material

**1. Supplementary Table S1.** Accession information for all samples included in this study.
**2. Supplementary Table S2.** Voucher specimens examined in this study.
**3. Supplementary File S1.** Circos plots of chloroplast genome assemblies for 49 samples of *Aristolochia* and related taxa.
**4. Supplementary File S2.** Species tree inferred using ASTRAL-III.
**5. Supplementary File S3.** Concatenated nuclear supermatrix tree.
**6. Supplementary File S4.** Plastid supermatrix phylogeny.
**7. Supplementary File S5.** Results from PhyloNet hybridization analysis.
**8. Supplementary File S6.** Results from PhyParts concordance analysis.
**9. Supplementary File S7.** Ancestral state reconstruction for RICH syndrome traits.

## Acknowledgements

We thank Dr. Israel Lopes da Cunha Neto, Dr. Lena Hunt, Mr. Zachary Kozma, Ms. Hannah Ratcliff, Ms. Angelique Acevedo, Ms. Annabelle Wang, Mr. Leo Semana, and Ms. Caterina Gandolfi (Onyenedum Lab, New York University, New York, USA) for continuous technical and botanical support throughout this project. We are also deeply grateful to Dr. Xinxin Zhu (Xinyang Normal University, Xinyang, China), Dr. Yunjuan Zuo and Mr. Xingchi Xie (Xishuangbanna Tropical Botanical Garden, Xishuangbanna, China), Dr. Matthew Pace and Dr. Dennis Stevenson (New York Botanical Garden, New York, USA), Dr. Pedro Acevedo-Rodríguez (Smithsonian Institution, Washington, DC, USA), Dr. Lei Cai, Dr. Jie Cai, and Mr. Jidong Ya (Kunming Institute of Botany, Kunming, China), Mr. Yiwen Jiang (Beihai, China), Dr. Sven Landrein (Kadoorie Farm and Botanic Garden, Hong Kong, China), Mr. Wesley Franks (Austin, USA), and Dr. Bin Liu and Dr. Lumei Liu (Institute of Botany, Chinese Academy of Sciences, Beijing, China), Mr. Yifan Li and Mr. Zhengxu Ma (Bazi Collection & Botanical Garden, Mengzi, China), and Mr. Dominik Frank (University of Tübingen, Tübingen, Germany), for valuable consultation, material exchange, and assistance with voucher deposit. We are especially grateful to Ms. Yushan Cai (Dingxi, China) for the illustrations of root systems and plant habit in Figure 1, and to Ms. Binyu Zhao (Suzhou University, Suzhou, China) for the illustrations in Figure 5, including panel (e) and panels (f–i). We thank Mr. Qing Yu (Ningbo, China) for silica sample collection, maintenance of living specimens, and participation in fieldwork. We also acknowledge the assistance of Ms. Lihua Zi (Kunming, China), Ms. Jing Li (Baoshan, China), Ms. Ying Yang, Ms. Tianping Huang, Ms. Han Lai, and Ms. Guijuan Wang (Xishuangbanna Tropical Botanical Garden, Xishuangbanna, China), Ms. Jihong Li and Ms. Xiao Li (Hong Kong, China), and Mr. Fuguan Deng and Mr. Mingfeng Long (Wenshan, China), for travel support and fieldwork facilitation.

Figure contributions are gratefully acknowledged as follows: Figure 1 — panel (h) by Dr. Boka Li (Beijing, China); panel (i) by Dr. Xinxin Zhu; panels (j) and (o) by Mr. Lianjie Li (Bengbu, China); panel (l) by Mr. Xinjie Zhao (Mengzi, China); and panel (q) by Dr. Sven Landrein. Figure 7 — panels (a) and (i–m) by Mr. Yiwen Jiang.

## Funding Source

This project was fully supported by a grant from the Center for Environmental and Animal Protection (CEAP) at NYU to Y.F. Wang. Additional support was provided in part by startup funds and a CAREER award to Dr. J.G. Onyenedum (NSF 2401675), the National Natural Science Foundation of China (32300178), a China Postdoctoral Science Foundation fellowship to Dr. S. Liao (2024M753278), and the Key Technology Research and Development Program of Zhejiang Province to Dr. P. Li (2023C03138). Computational support was provided in part by NYU IT High Performance Computing resources and services.

---

**Figure S1.**
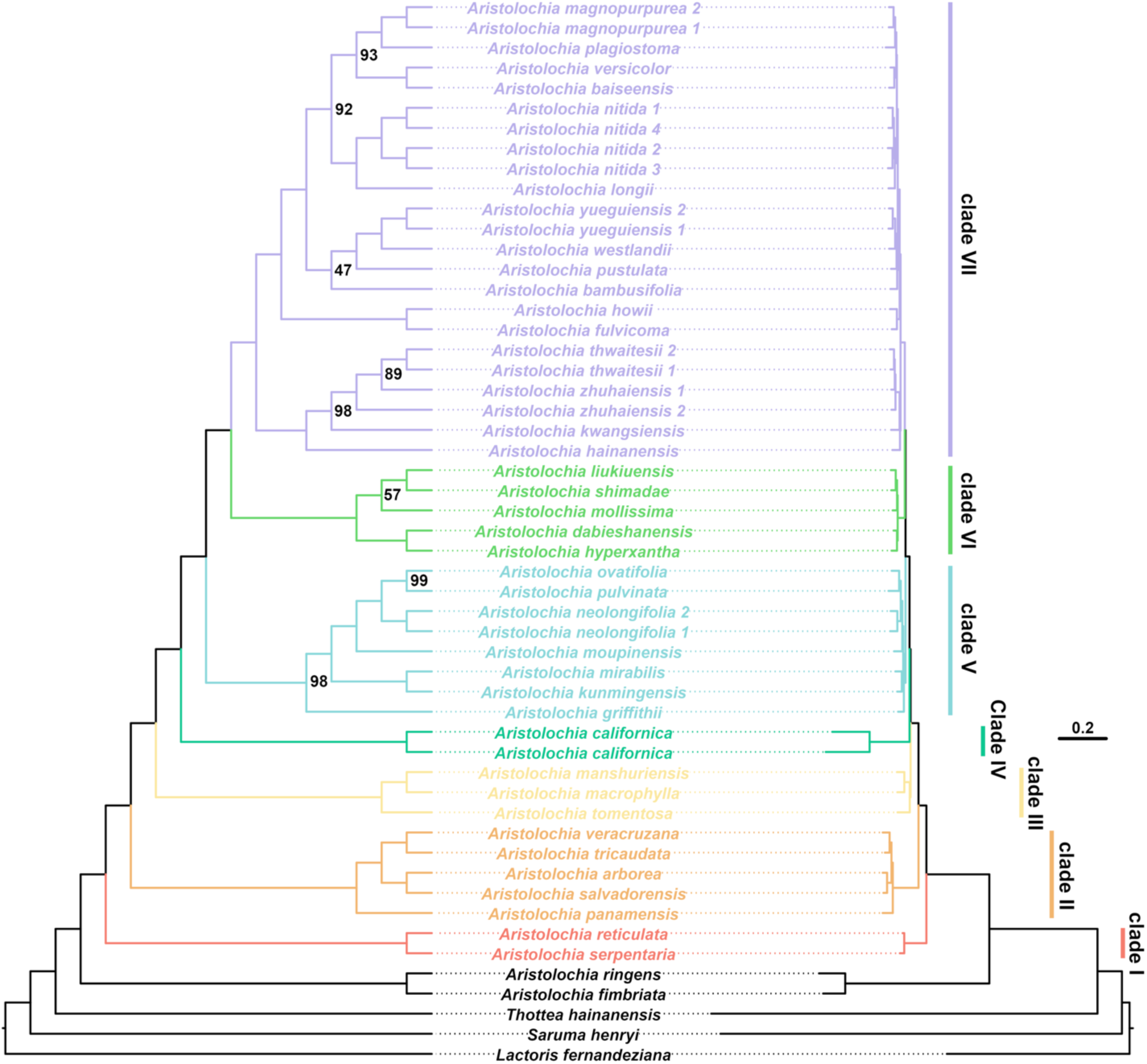
Nuclear phylogeny inferred from Angiosperms353 concatenated (“supermatrix”) data. The topology on the left is shown in rectangular layout without branch lengths, with bootstrap support values indicated at nodes; unlabeled nodes denote full support (BS = 100). The right panel presents the same topology with branch lengths illustrated. Clade color coding is consistent with clade delineations presented in Figures 1–3 and throughout the manuscript.

**Figure S2.**
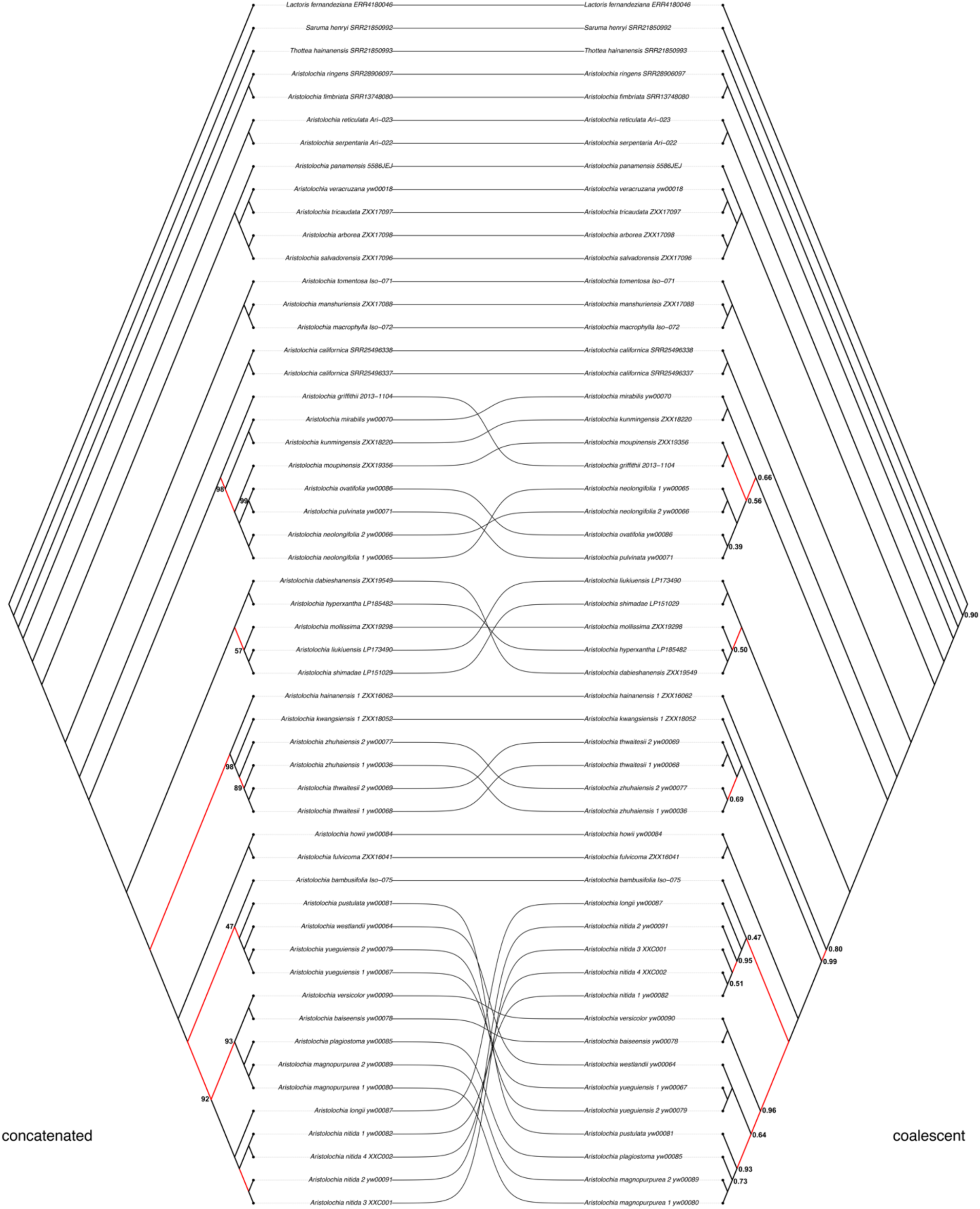
Cophylogeny comparison of nuclear topologies inferred using two methods. The left panel shows the concatenated (supermatrix) phylogeny; the right panel presents the coalescent-based species tree (ASTRAL-III). Nodes without labeled support values indicate full support (BS = 100 for concatenated; LPP = 1.0 for coalescent). Branches highlighted in red indicate topological conflicts between the two trees.

**Figure S3.**
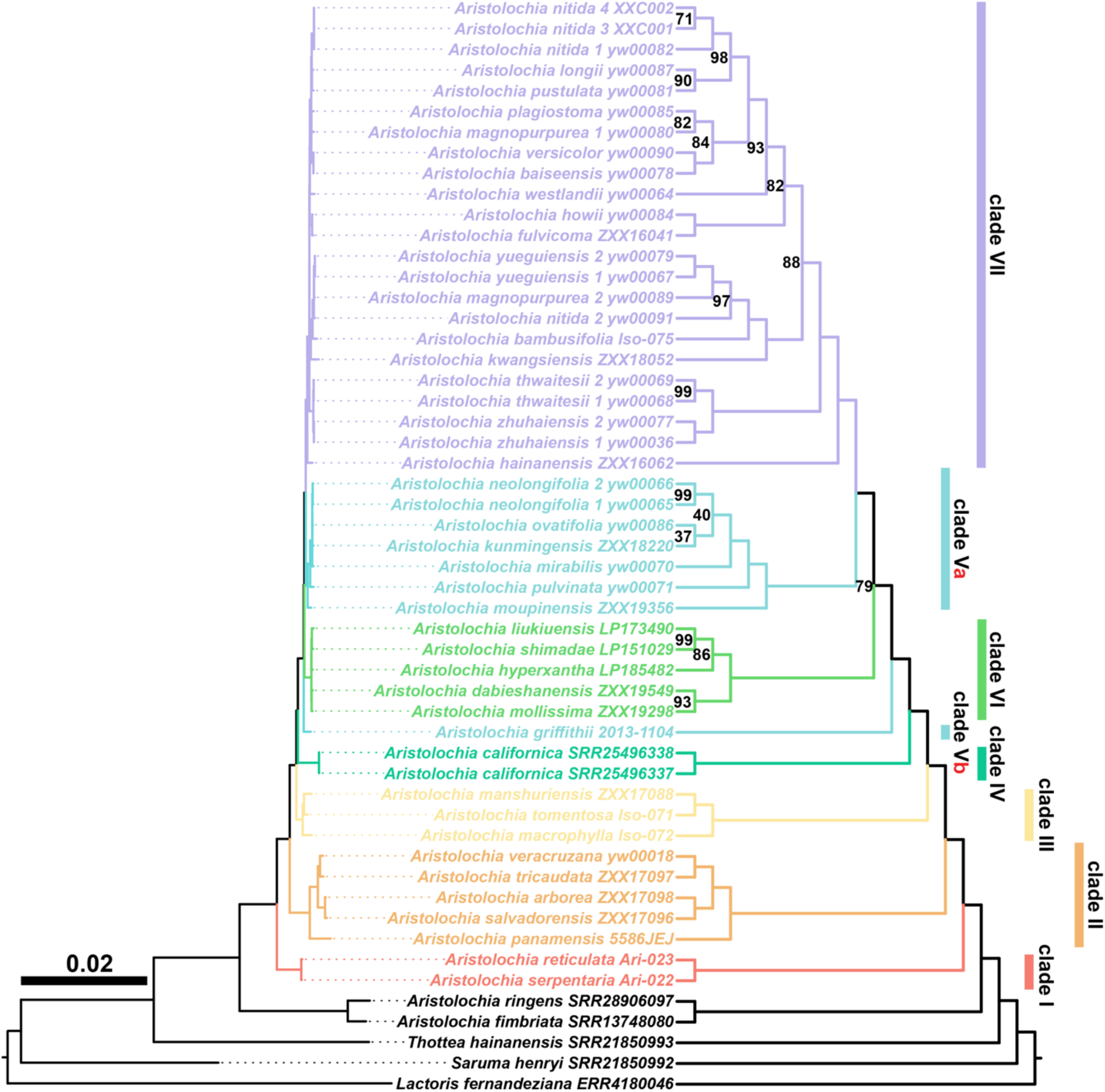
Chloroplast phylogeny inferred via maximum likelihood from complete plastome coding sequences (CDS), including tRNA, rRNA, and open reading frames. The left panel depicts the topology with branch lengths, while the right panel shows the same topology in rectangular format with bootstrap support values indicated at nodes; unlabeled nodes received full support (BS = 100). Clade color coding follows that used in Figures 1–3 and Figure S1, and is consistent throughout the manuscript. Notably, accession 2014-2014 (*Aristolochia griffithii*) is consistently placed in Clade V in previous analyses, but in this chloroplast tree it is not recovered as monophyletic with the rest of Clade V taxa. Accordingly, the original clade is denoted as Clade Va, and *A. griffithii* is treated separately as Clade Vb.

**Figure S4.**
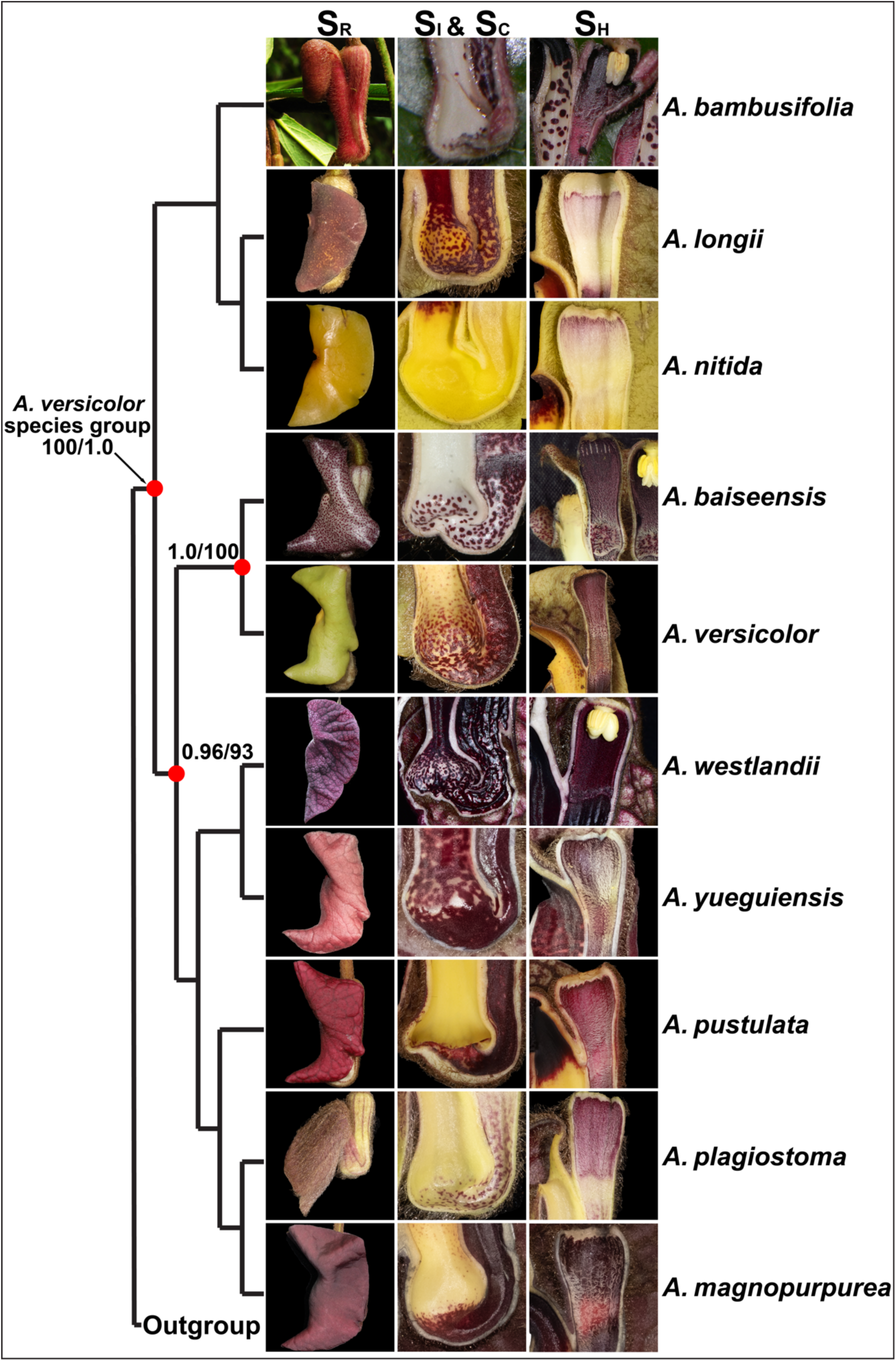
Illustration of the “RICH” syndrome in the *Aristolochia versicolor* species group. The left panel shows the species tree topology for this clade. Diagnostic floral synapomorphies are annotated as follows: **SR**: reflexed calyx limb; **SI** (left tube): inflated upper tube; **SC** (right tube): constricted basal tube entrance; **SH**: hairy band on the adaxial side of the utricle, distinctly two-tiered and surrounding the gynostemium. The reflexed calyx (**SR**) is absent in *A. plagiostoma* and *A. bambusifolia*, while the remaining synapomorphies are consistently present across all members of the clade and absent from other *Siphisia* species.

## Notes

### Competing Interest Statement

The authors have declared no competing interest.

